# TRIM9 switches the morphological phenotype of melanoma cells

**DOI:** 10.64898/2026.03.17.712420

**Authors:** Kimberly Lukasik, Aneri B. Shah, Chris T. Ho, Matthew Li, G. Becky Patrick, Jordan Brooks, Simon Rothenfusser, James E. Bear, Stephanie L. Gupton

## Abstract

Melanoma is a highly plastic cancer characterized by distinct cellular phenotypes associated with broadly unique gene expression profiles. TRIM9 is a brain-enriched E3 ubiquitin ligase detected in melanoma, but how TRIM9 expression regulates melanoma phenotype is unknown. Using two metastatic human melanoma cell lines and a mouse melanoma model, we found that TRIM9 promoted melanoma proliferation and altered cell morphology. In cell lines, TRIM9 promoted cellular blebbing and negatively regulated adhesion, secretion, and mesenchymal motility. TRIM9 interacted with VASP in melanoma cells, altering VASP modification, localization, and dynamics. In the absence of TRIM9, cells had an altered actin organization and more focal adhesions, where VASP accumulated and exhibited rapid turnover. We find the alterations in actin architecture and adhesion associated with TRIM9 deletion were coincident with increased motile and contractile mesenchymal behavior in vitro. In vivo loss of TRIM9 in melanoma slowed tumor growth and altered metastasis frequency, size, and destination. Our findings indicate TRIM9 alters the proliferative and morphological phenotypes of metastatic melanoma cells to influence disease progression.

## Introduction

Melanoma is a particularly plastic cancer, in which cells undergo phenotype switching, with distinct cellular states associated with broadly unique gene expression profiles and proliferative phenotypes (Widmer et al., 2012; Rambow et al., 2019; Hossain and Eccles, 2023; Wouters et al., 2020; Andrews et al., 2022; Pozniak et al., 2024). These include proliferative melanocytic states, invasive neural-crest like states, mesenchymal states, and one or more transitional states. The heterogeneity of these states is core to the aggressive nature of melanoma, including the ability to respond to nutrient deprivation and gain resistance to immune checkpoint inhibitors.

Defining the differentially expressed genes (DEGs) that influence melanoma phenotype switching is critical. A recent study surveyed gene expression across > 60 human melanoma lines derived from patient samples (Kutt et al., 2021). Machine learning identified the 15 highest ranked DEGs among the 19,000 genes considered that could distinguish clusters of melanoma cell lines. Of these 15 genes, 3 genes: GSTO1, TBC1D16, and TRIM9 were able to reliably classify nearly all lines (Kutt et al., 2021). TRIM9 piqued our interest, as it is a brain-enriched E3 ubiquitin ligase we had characterized as a potent regulator of neuronal morphogenesis (Menon et al., 2015; Plooster et al., 2017; McCormick et al., 2024a; Mutalik et al., 2025). TRIM9 is frequently detected in melanoma biopsies (Pozniak et al., 2024; Kutt et al., 2021; TCGA Research Network; Vilaseca et al., 2023; Bartley et al., 2023), but is not expressed in melanocytes (Bartley et al., 2023; Hodis et al., 2022). Melanocytes derive from the neural crest lineage. Mining single cell RNA sequencing (scRNAseq) studies revealed that TRIM9 is expressed broadly in the neural crest lineage during development (Soldatov et al., 2019; Pijuan-Sala et al., 2019). Analogously, scRNAseq studies of patient-derived melanomas indicate that TRIM9 expression is high in melanoma cells occupying an intermediate or neural crest-like state (Wouters et al., 2020; Andrews et al., 2022; Pozniak et al., 2024). This dedifferentiated neural crest-like state often exhibits stemcell–like properties and is associated with impaired immune recognition and resistance to T-cell-mediated therapies (Pozniak et al., 2024; Restivo et al., 2017; Civenni et al., 2011; Mehta et al., 2018; Boiko et al., 2010; Landsberg et al., 2012). However, how TRIM9 alters melanoma phenotype is not known.

TRIM9 is a brain-enriched member of the tripartite motif containing (TRIM) family of Really Interesting New Gene (RING) E3 ubiquitin ligases (Berti et al., 2002). Although detected at low levels outside the nervous system, several cancers demonstrate high TRIM9 expression (TCGA Research Network). TRIM9 suppresses tumor growth in pancreatic cancer and glioblastoma, and suppresses migration and invasion in pancreatic and esophageal cancers (Liu et al., 2018; Lin et al., 2023). In breast cancer, TRIM9 promoter methylation and epigenetic silencing was a suggested tumor suppressor (Mishima et al., 2015). In contrast, TRIM9 promoted tumor progression and motility in bladder cancer (Zhang et al., 2023) and non-small cell lung cancer (Wang et al., 2023). Thus, TRIM9 regulates core processes including motility and growth, but its impact on tumor progression is context-dependent.

From our work in neurons, we know that TRIM9 interacts with the netrin receptors DCC and UNC5 and regulates morphological responses to the extracellular guidance cue netrin-1 (Winkle et al., 2014; Menon et al., 2015; Winkle et al., 2016; Plooster et al., 2017; Urbina et al., 2018; Menon et al., 2020, 2021; McCormick et al., 2024a; Mutalik et al., 2025). TRIM9 interacts with the actin polymerase VASP at filopodial tips and negatively regulates its function via non-degradative ubiquitylation (Menon et al., 2015; McCormick et al., 2024b). TRIM9 interacts with and negatively regulates the secretory t-SNARE, SNAP25 (Winkle et al., 2014; Urbina et al., 2018). Murine *Trim9* deletion increases filopodial stability and secretion, causes excessive axonal and dendritic branching, and impairs netrin-dependent axon firing and netrin-dependent axon turning (Winkle et al., 2014; Menon et al., 2015; Plooster et al., 2017; Urbina et al., 2018; McCormick et al., 2024a; Mutalik et al., 2025). These interactions indicate that TRIM9 may also be poised to alter morphology in melanoma.

Notably, the TRIM9 substrate VASP is implicated in pro-oncogenic processes in multiple cancers, via regulation of cytoskeletal remodeling and cell migration (Han et al., 2008; Xiang et al., 2018; Liu et al., 2019; Li et al., 2020; Gui et al., 2024). VASP localizes to the tips of bundled actin-based filopodial protrusions, the leading edge of fan-like actin-based lamellipodial protrusions, and focal adhesions, large multi-layered protein complexes responsible for adherence of cells to the extracellular environment (Reinhard et al., 1992; Bear et al., 2002; Lebrand et al., 2004). VASP accelerates actin filament elongation, promotes filament bundling, and protects the growing filament end from capping proteins (Barzik et al., 2005; Breitsprecher et al., 2008). Within the actin-regulatory layer of focal adhesions, VASP promotes actin filament polymerization and bundling, coupling the adhesion to the actin cytoskeleton (Chung et al., 2025). Whether TRIM9-dependent regulation of VASP may influence cancer cell behavior is not known.

To address the role of TRIM9 in melanoma, we utilized two metastatic human melanoma cell lines and a genetically engineered mouse in which we could induce endogenous melanoma on the ear. In each model, we defined consequences of TRIM9 loss, finding that TRIM9 was a potent regulator of melanoma cell morphology, proliferation, and migratory behavior. Using cell lines, we show that TRIM9 loss inhibited blebbing and increased cell spreading, altered filamentous actin architecture and increased focal adhesions. We found that TRIM9 interacted with VASP, altering VASP phosphorylation and localization in the cell. In the absence of TRIM9, VASP demonstrated increased localization to focal adhesions, where it exhibited accelerated dynamics. We found that these alterations in actin architecture and adhesion associated with TRIM9 loss were coincident with increased mesenchymal motility, secretion, and invasion in vitro. In vivo, loss of TRIM9 slowed tumor progression and altered the frequency, size, and location of metastases. Together our findings indicated that TRIM9 expression promotes proliferative, morphological, and migratory phenotype switching of metastatic melanoma to promote tumor progression.

## Results

### TRIM9 is expressed in human melanoma and promotes blebbing

TRIM9 mRNA and protein are detectable in human melanoma (Bartley et al., 2023; Vilaseca et al., 2023; Andrews et al., 2022; Wouters et al., 2020; Pozniak et al., 2024; Arafeh et al., 2025; DepMap, Broad, 2025; TCGA Research Network). High TRIM9 mRNA expression was associated with reduced survival in both untreated (hazard ratio HR =1.6, log rank *P*=9*e*^−04^, n =397 patients, **Fig S1A**) and immune checkpoint inhibitor treated patients (anti-PD-1, HR=1.42 log rank *P* = 0.029, n = 334 patients, anti-CTLA-4 HR = 1.93, log rank *P* = 0.016, n = 112 patients,(**Fig S1B-C**) (Kovács et al., 2023). Since high TRIM9 RNA was hazardous, we evaluated TRIM9 protein in two metastatic human melanoma lines with high TRIM9 mRNA levels, 1205Lu and WM266-4 (Arafeh et al., 2025; DepMap, Broad, 2025; Juson-Torres, 2025). Immunoblotting revealed TRIM9 long and short isoforms were present in both lines. To investigate the role of TRIM9, we utilized CRISPR-Cas9 genome editing to delete both TRIM9 isoforms and observed total loss of detectable protein (**Fig 1A**). In developing neurons, we found that TRIM9 was a potent regulator of neuronal morphology (Winkle et al., 2014; Menon et al., 2015; Plooster et al., 2017; McCormick et al., 2024a). We first aimed to determine whether melanoma cell morphology was similarly altered by loss of TRIM9. Loss of TRIM9 was associated with an increased cell area and a less circular, more polarized morphology in 1205Lu cells (**Fig 1B-D**) and WM266-4cells (**Fig S1D-F**). We also found that WM266-4 *TRIM9*^*+/+*^ cells exhibited increased number of cells with bleb-like morphology on glass (**Fig S1G**); this bleb-like morphology was not observed in 1205Lu cells on glass.

**Fig. 1.**
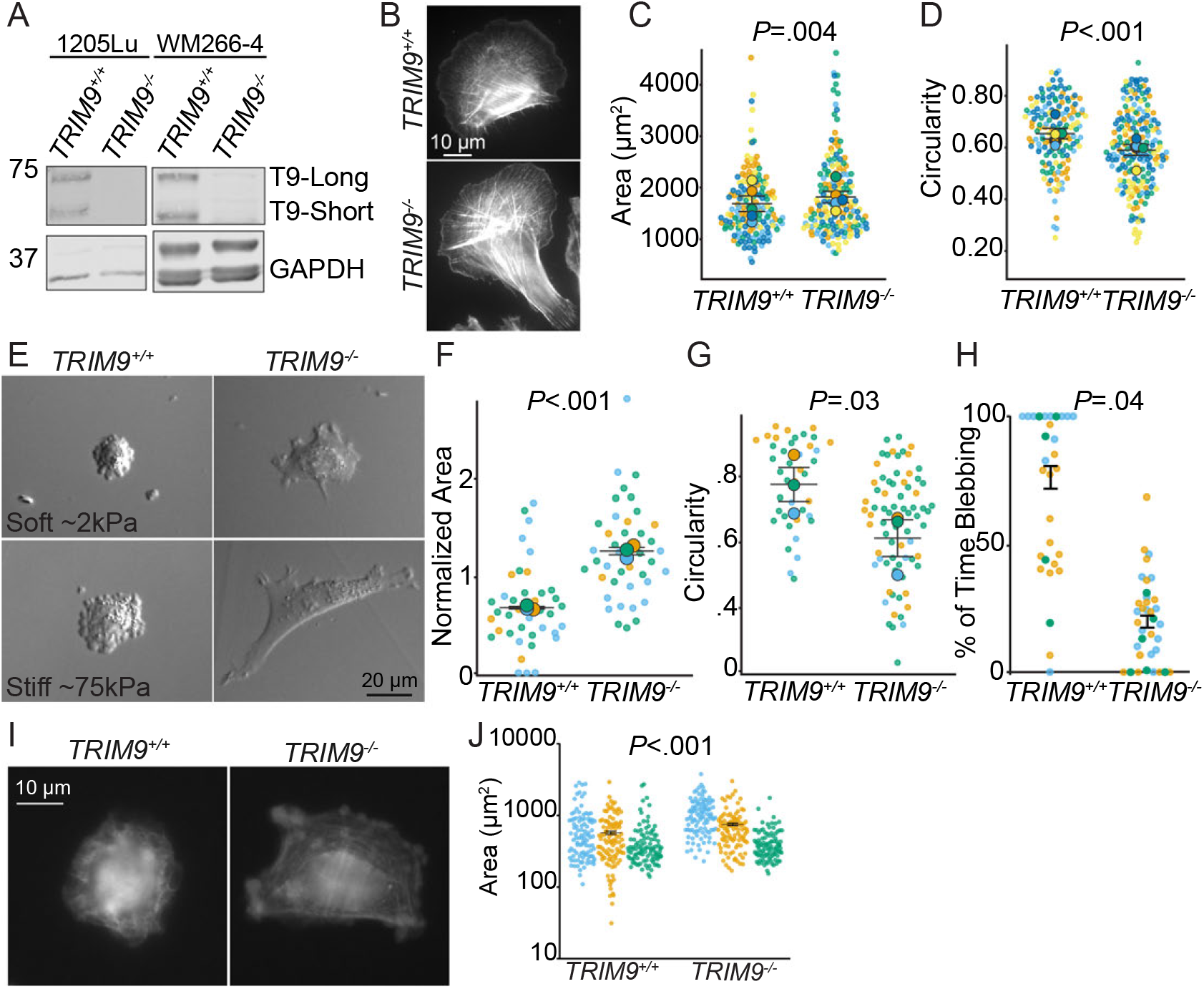
TRIM9 is expressed in human melanoma and alters cell morphology. (**A**) Representative western blot of 1205Lu and WM266-4 lysates. GAPDH is used as a loading control. CRISPR/Cas9 generated *TRIM9*^−/−^ cells lack both short and long TRIM9 isoforms. (**B**) Representative images of 1205Lu *TRIM9*^*+/+*^ and *TRIM9*^−/−^ cells plated on glass coverslips and stained for F-actin (phalloidin). (**C**) Cell area of 1205Lu cells on glass, *TRIM9*^−/−^ cells were larger than *TRIM9*^*+/+*^ cells (*P*=0.004, two-tailed *t*-test on log transformed data), n = 5 independent biological replicates (unique colors); n ≥ 20 cells per genotype per replicate, each dot indicates a cell. (**D**) Cell circularity on glass, *TRIM9*^*+/+*^ cells were more circular (circle =1.0) (*P* <.001, two-tailed *t*-test), n = 5 independent replicates; n ≥ 20 cells per genotype per replicate, each dot represents a cell. (**E**) 1205Lu cells plated on soft (~ 2 kPa) and stiff (~ 75 kPa) polyacrylamide hydrogels. (**F**) 1205Lu cell area on soft hydrogels, loss of TRIM9 increased cell area (*P* <.001, two-tailed *t*-test), n = 3 independent biological replicates; n ≥ 39 cells per genotype, each dot indicates a cell. (**G**) Cell circularity on soft hydrogels; *TRIM9*^*+/+*^ cells are more circular than *TRIM9*^−/−^ cells (*P* =0.03, two-tailed *t*-test), n = 3 independent biological replicates-unique colors; n ≥ 39 cells per genotype per replicate, each dot indicates a cell. (**H**) Time spent in a bleb-like state on soft hydrogels; *TRIM9*^*+/+*^ cells spend more time blebbing (*P* =.04, two-tailed *t*-test on log transformed data), n = 3 independent biological replicates; n ≥ 39 cells per genotype, each dot indicates a cell. (**I**) Phalloidin stained 1205Lu cells plated on glass for two hours. (**J**) Cell area for cell spreading; *TRIM9*^−/−^ cells exhibit increased spreading at two hours as compared to *TRIM9*^*+/+*^ (*P* <.001, two-tailed *t*-test on natural log transformed data), n = 3 independent biological replicates; n ≥ 100 cells per genotype per replicate, each dot represents a cell.

Melanoma cells often adopt a bleb-like morphology for migration and cell survival in low attachment (Cantelli et al., 2015; Gabbireddy et al., 2021; Weems et al., 2023; Driscoll et al., 2024). Blebbing is a consequence of the detachment of cortical actin from the plasma membrane and decreased cell adhesion, and is noted by a more circular, less spread state (Tinevez et al., 2009; Paluch et al., 2006). Substrate stiffness alters melanoma phenotypes, including morphology (Diazzi et al., 2023; Lecacheur et al., 2024). To determine whether TRIM9 modulated melanoma cell morphology on substrates with physiologically relevant stiffnesses, we plated 1205Lu cells on soft (~ 2 kPa) and stiffer (~ 75 kPa) polyacrylamide hydrogels (**Fig 1E**). Although 1205Lu cells did not bleb on glass, both *TRIM9*^*+/+*^ and *TRIM9*^−/−^ 1205Lu cells transitioned from flat, mesenchymal like states to bleb-like morphological states on hydrogels (**Fig 1E**). 2 kPa mimics the soft extracellular matrix (ECM) of the dermal and early tumor environment as well as common metastatic sites such as the brain, whereas 75 kPa is a very stiff matrix, more associated with tumor tissues (Park et al., 2023; Staunton et al., 2016; Massey et al., 2024; Micalet et al., 2023; Wahlsten et al., 2023). Both *TRIM9*^*+/+*^ and *TRIM9*^−/−^ cells spread more on stiff hydrogels than soft hydrogels (**Fig S1H-I**). *TRIM9*^−/−^ cells exhibited an increased cellular area on both soft (**Fig 1F**) and stiff hydrogels (**Fig S1J**), indicating loss of TRIM9 induced a more spread phenotype, as on glass. We noted changes to cellular symmetry and polarization on hydrogels. Both *TRIM9*^*+/+*^ and *TRIM9*^−/−^ cells were more circular, less polarized on soft substrates (**Fig S1K-L**). On both soft and stiff hydrogels *TRIM9*^−/−^ cells were less circular than *TRIM9*^*+/+*^ cells (**Fig 1G**,**Fig S1M**). Like WM266-4 cells on glass, on the 2 kPa substrates specifically, *TRIM9*^*+/+*^ 1205Lu cells spent approximately 3.4x more time blebbing than *TRIM9*^−/−^ cells, (**Fig 1H, VideoS1**). We next investigated whether loss of TRIM9 affected the early stages of spreading (**Fig 1I**). TRIM9 loss resulted in increased cell spreading two hours after plating on glass (**Fig 1J**), with an increased distribution of larger cells (**Fig S1N**). Together these results indicated that in melanoma cells TRIM9 negatively influenced cell spreading and promoted blebbing and was thus a potent regulator of melanoma cell morphology.

### Loss of TRIM9 alters filamentous actin structures

Since TRIM9 loss in melanoma cells altered cell morphology, we next investigated how loss of TRIM9 altered the filamentous actin (F-actin) cytoskeleton. F-actin in melanoma cells is organized into stress fiber-like actin bundles, filopodia, and lamellipodia (**Fig 2A**). Total F-actin levels revealed by phalloidin staining were unchanged by TRIM9 loss (**Fig 2B**). In developing neurons, loss of TRIM9 increased filopodial length and density in the axonal growth cone, and growth cone size. We hypothesized that analogous actin-based protrusions in melanoma would be altered by loss of TRIM9. Some 1205Lu cells exhibited both filopodial and lamellipodial protrusions, whereas others did not have filopodia (**Fig 2C**,**Fig S2A**). Unlike neurons, loss of TRIM9 in 1205Lu cells reduced filopodial length (**Fig 2D**) and density along the leading edge (**Fig 2E**), but increased filopodia width (**Fig 2F**). We next examined lamellipodia. Loss of TRIM9 in melanoma resulted in decreased width of the lamellipodia (**Fig 2G**). However, the length of the lamellipodia along the cell periphery was unchanged (**Fig S2B**). There was no difference in F-actin intensity within lamellipodia (**Fig S2C**).

**Fig. 2.**
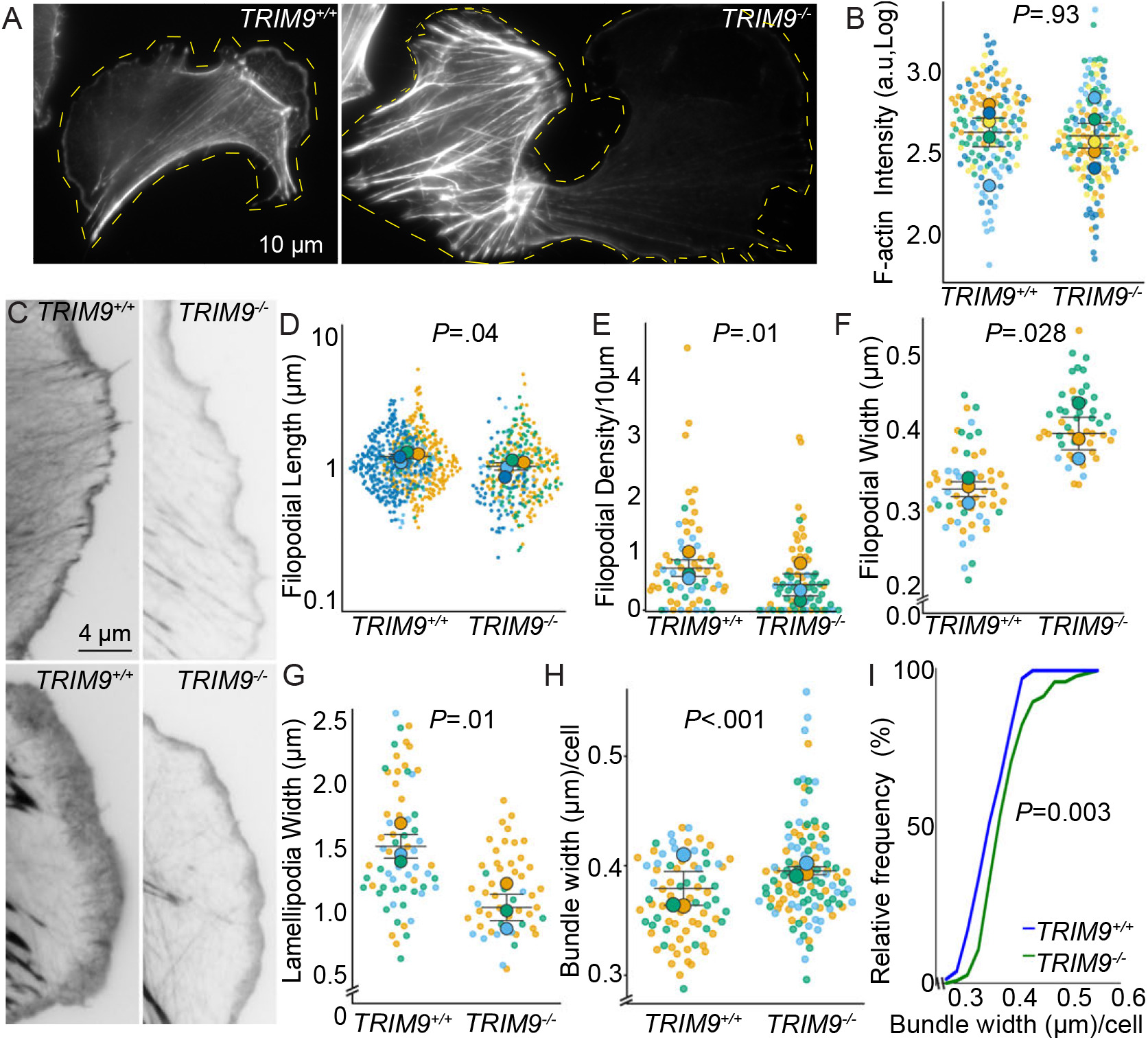
Loss of TRIM9 alters F-actin organization. (**A**) Representative images of 1205Lu *TRIM9*^*+/+*^ and *TRIM9*^−/−^ cells labeled for phalloidin (F-actin). (**B**) Cellular phalloidin intensity of 1205Lu *TRIM9*^*+/+*^ and *TRIM9*^−/−^ cells are not different (*P* =0.93, two-tailed *t*-test), n = 5 independent biological replicates; n ≥ 20 cells per genotype per replicate. (**C**) Images of the leading edge of phalloidin-stained 1205Lu cells. (**D**) Filopodial length in 1205Lu cells; *TRIM9*^−/−^ cells had reduced filopodial length (*P* =0.04, two-tailed *t*-test on log transformed data), n = 4 independent biological replicate, each color represents a separate experiment; n ≥ 10 cells per genotype per replicate, each dot represents individual filopodium. (**E**) Filopodial density/10 µm of cell perimeter in 1205Lu cells; *TRIM9*^−/−^ cells had reduced filopodial density (*P* =0.01, two-tailed *t*-test), n = independent biological replicates, n ≥ 10 cells per genotype per replicate, each dot represents the average filopodial density per cell. (**F**) Average filopodia width per cell; TRIM9 loss increased filopodial width (*P* =.028, two-tailed *t*-test), n = 3 independent biological replicates, n ≥ 7 cells per genotype per replicate, each dot represents the average filopodial width per cell. (**G**) Lamellipodial width; TRIM9 loss reduced lamellipodial width (*P* =.01, two-tailed *t*-test), n = 3 independent biological replicates, n ≥ 7 cells per genotype per replicate, each dot represents the average lamellipodium width per cell. (**H**) Actin stress fiber bundle width; loss of TRIM9 increased average bundle width (*P* <.001, Mann-Whitney test), n = 3 independent biological replicates, n ≥78 cells per genotype, each dot is cell. (**I**) The cumulative distribution of bundle width was shifted to the right in *TRIM9*^−/−^ cells (*P* <.001, Kolmogorov-Smirnov test (KS)).

We next evaluated stress fiber organization. We found that loss of TRIM9 increased the average stress fiber bundle width per cell (**Fig 2H**). Cumulative distribution analysis of the average stress fiber width per cell revealed a rightward shift associated with loss of TRIM9, corresponding to an increased bundle thickness (**Fig 2I**). Similar results were observed when analyzing individual fibers (**Fig S2D-E**). These data suggested that TRIM9 modulated multiple F-actin structures in human melanoma.

### TRIM9 alters VASP levels in filopodia and lamellipodia

We were curious why loss of TRIM9 resulted in distinct filopodial and lamellipodial phenotypes in melanoma and neurons. Ena/VASP proteins are actin polymerases that localize to both lamellipodial and filopodial actin structures in fibroblasts and neurons (Lanier et al., 1999; Bear et al., 2002; Lebrand et al., 2004). We previously found that TRIM9 interacted with and modified the actin polymerase VASP with multi-mono-ubiquitin, a post translational modification (PTM) that altered actin polymerase function, and that loss of TRIM9 increased VASP localization to filopodia tips in neurons (Menon et al., 2015; Boyer et al., 2020; McCormick et al., 2024b). TRIM9 also localized to filopodial tips and to the leading edge of lamellipodia in melanoma (**Video S2**). To determine whether loss of TRIM9 regulated VASP in melanoma cells, we immuno-labeled VASP. In *TRIM9*^*+/+*^ 1205Lu cells, VASP immunolocalization highlighted filopodia tips, lamellipodia tips, and dispersed punctate structures behind the cell edge (**Fig 3A**). In contrast in *TRIM9*^−/−^ cells, VASP localized similarly at the cell edge but enhanced dramatically behind the cell edge. Quantification revealed that loss of TRIM9 increased total VASP levels in the cell (**Fig 3B**), with the percentage of cellular VASP localized to the lamellipodia and to filopodia tips decreased (**Fig 3C-D**). This reduction was due to cellular increases in VASP, as comparison of absolute VASP immunostaining at lamellipodia and filopodia was increased in *TRIM9*^−/−^ cells (**Fig 3E-F**). In *TRIM9*^*+/+*^ cells, VASP was enriched at the tips of filopodia and not often detected behind the tip (**Fig 3A insets, 3G**). In contrast, in addition to tip enrichment, VASP often appeared in multiple puncta along filopodia in *TRIM9*^−/−^ cells (**Fig 3A insets, 3H**). To reveal this difference in VASP distribution, we normalized F-actin and VASP fluorescence intensity across an even number of bins along filopodia lengths (**Fig 3I**). This demonstrated VASP tip enrichment in *TRIM9*^*+/+*^ cells and a distinct distribution along filopodia in *TRIM9*^−/−^ cells. Taken together these data suggested that TRIM9 regulated both branched and bundled F-actin protrusions in melanoma cells and altered VASP localization to these actin structures.

**Fig. 3.**
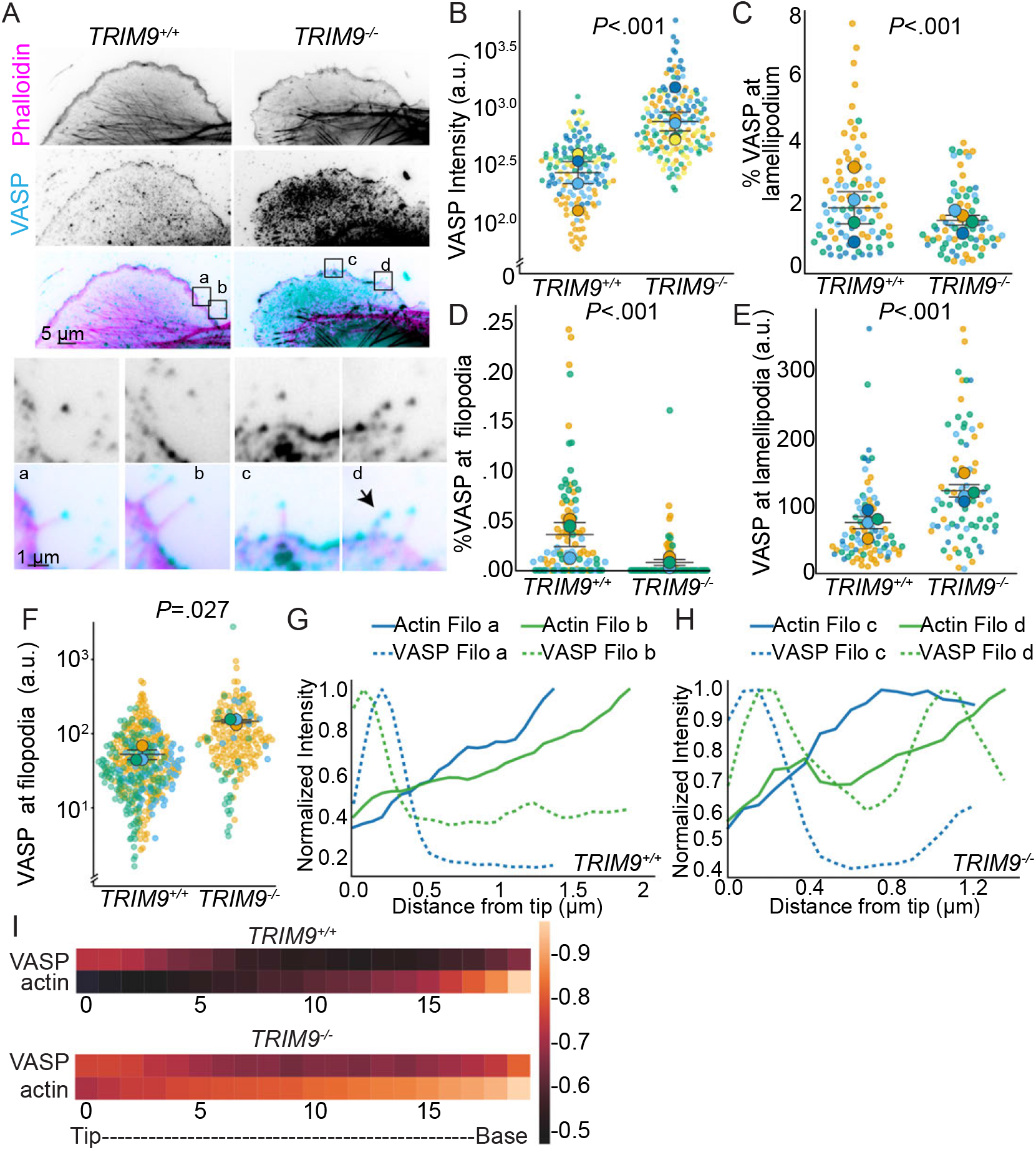
Loss of TRIM9 alters VASP localization to actin-based structures. (**A**) Representative images of fixed 1205Lu cells labeled for phalloidin (F-actin) and VASP with corresponding labeled insets. (**B**) Total cell levels of VASP were increased in *TRIM9*^−/−^ cells (*P* <.001, two-tailed *t*-test on log transformed data), n = 5 independent biological replicates, each color is a replicate; n ≥ 20 cells per genotype per replicate, each dot is a cell. (**C**) The percentage of VASP localized to lamellipodium was reduced by loss of TRIM9 (*P* <.001, Mann-Whitney test), n = 4 independent biological replicates, n ≥ 10 cells per genotype per replicate, each dot is a cell. (**D**) Loss of TRIM9 reduced the percentage of total cellular VASP at filopodia (*P* <.001, Mann-Whitney test), n = 3 independent biological replicates, n ≥ 16 cells per genotype per replicate, each dot is a cell. (**E**) Loss of TRIM9 increased the intensity of VASP at lamellipodia (p<.001, Mann-Whitney test on log transformed data), n = 4 independent biological replicates, n ≥ 11 cells per genotype per replicate, each dot is a cell. (**F**) Loss of TRIM9 increased the intensity of VASP at filopodial tips (*P* =.027, Mann-Whitney test), n = 3 independent biological replicates, n ≥ 16 cells per genotype per replicate. (**G-H**) Line scans of filopodial VASP and phalloidin 1205Lu cells, inserts from A, indicate alteration in localization of VASP across filopodia associated with loss of TRIM9. (**I**) Heat map of VASP localization. Linescans were drawn from the tip (0) to the base of the filopodial and represented as a heatmap localization of VASP throughout the length of the filopodia. To generate the heatmap, fluorescence intensity of each protein was normalized and length binned. n = 3 independent biological replicates; n ≥ 266 filopodia per genotype.

### Loss of TRIM9 perturbs VASP localization and focal adhesion number and shape

We next evaluated the enhanced VASP behind the cell edge in cells lacking TRIM9 (**Fig 3A**). VASP in both human and mouse melanoma cells, fibroblasts, and HEK cells also localizes to focal adhesions (Reinhard et al., 1992; Benz et al., 2009; Döppler et al., 2013; McCormick et al., 2024b; Damiano-Guercio et al., 2020; Cheng and Mullins, 2020). Focal adhesions link actin bundles to ECM receptors at the plasma membrane. We observed transient TRIM9 localization to VASP-containing focal adhesions in 1205Lu cells (**Video S2**). To determine whether VASP localization at focal adhesions was altered by loss of TRIM9, we immuno-labeled VASP and the focal adhesion component paxillin, and stained F-actin with phalloidin in 1205Lu cells (**Fig 4A**,**Fig S4A**) and WM266-4 cells (**Fig S4C**). Again, higher VASP levels were observed behind the lamellipodia in cells lacking *TRIM9*, as noted when VASP fluorescence was scaled identically (**Fig 4A**). Boosting image contrast in *TRIM9*^*+/+*^ cells revealed that both VASP and paxillin localized to focal adhesions (**Fig S4A**). The percentage of cellular VASP localized to paxillin-positive focal adhesions increased in *TRIM9*^−/−^ 1205Lu cells (**Fig 4B**). This contrasted with VASP localization at lamellipodia and filopodia (**Fig 3C-D**). There was a higher absolute concentration of VASP at paxillin-positive focal adhesions (**Fig 4C**). TRIM9 loss in 1205Lu cells was also associated with increased cellular paxillin levels (**Fig 4D**). TRIM9 loss also increased the number of focal adhesions per cell in 1205Lu cells (**Fig 4E**) and WM266-4 cells (**Fig S4C-E**), and these focal adhesions were longer and narrower (**Fig 4F, Fig S4F**). Together these data indicated TRIM9 regulated VASP localization to focal adhesions and focal adhesion shape and number.

**Fig. 4.**
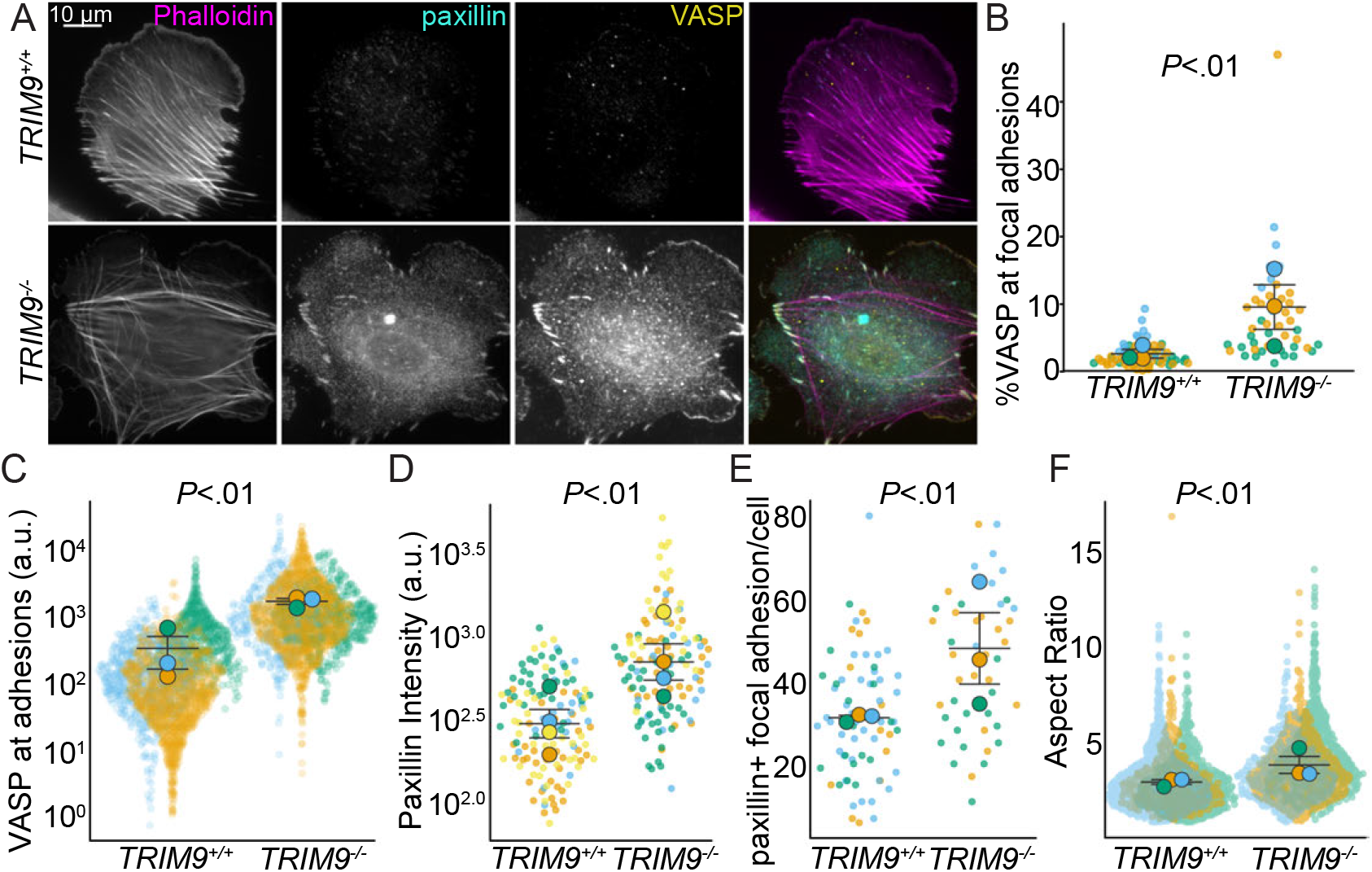
Loss of TRIM9altersVASP localization to focal adhesions, focal adhesion number, and morphology. (**A**) 1205Lu cells stained for phalloidin (F-actin), VASP, and paxillin. (**B**) Percentage of cellular VASP at paxillin-positive focal adhesions; loss of TRIM9 increased the percentage of cellular VASP localized to adhesions (*P*<.01, Mann-Whitney test), n = 3 independent biological replicates, each color is a unique replicate; n 7 cells per genotype per replicate, each dot represents a cell. (**C**) VASP intensity at focal adhesions; loss of TRIM9 increased VASP intensity (*P* <.01, Mann-Whitney test), n = 3 independent biological replicates; n 395 focal adhesions per replicate, each dot represents an adhesion. (**D**) Cellular paxillin intensity; loss of TRIM9 increased paxillin levels (*P* <.01, two-tailed *t*-test), n = 4 independent biological replicates, n 21 cells per genotype per replicate, each dot is a cell. (**E**) Paxillin-positive focal adhesions per cell; loss of TRIM9 increased adhesion number (*P* <0.01, two-tailed *t*-test), n = 3 independent biological replicates; n ≥ 7 cells per genotype per replicate. (**F**) Quantification of aspect ratio (width: height ratio) of paxillin-positive focal adhesions; loss of TRIM9 increased focal adhesion aspect ratio (*P* <.01, two-tailed *t*-test on log transformed data), n = 3 independent biological replicates; n 395 focal adhesions per genotype per replicate, each dot represents a focal adhesion.

### TRIM9 interacts with VASP and alters VASP phosphorylation

We identified VASP as an interaction partner and substrate of TRIM9 in neurons and HEK293 cells (Menon et al., 2015; Boyer et al., 2020). To evaluate whether this relationship persisted in melanoma, we performed coimmunoprecipitation using a mutant of TRIM9 lacking the ligase domain (TRIM9ΔRING). Loss of the RING domain stabilizes the interaction between ligases and substrates (Menon et al., 2015). We transiently expressed Halo-tagged TRIM9ΔRING with either VASP-GFP or GFP in 1205Lu (**Fig 5A**) and WM266-4 cells (**Fig S5A**) and performed a GFP immunoprecipitation. TRIM9ΔRING was detected in the immunoprecipitate with VASP-GFP, but not GFP alone (**Fig 5A**,**Fig S5A**). This indicated that TRIM9 interaction with VASP was maintained in melanoma cells.

**Fig. 5.**
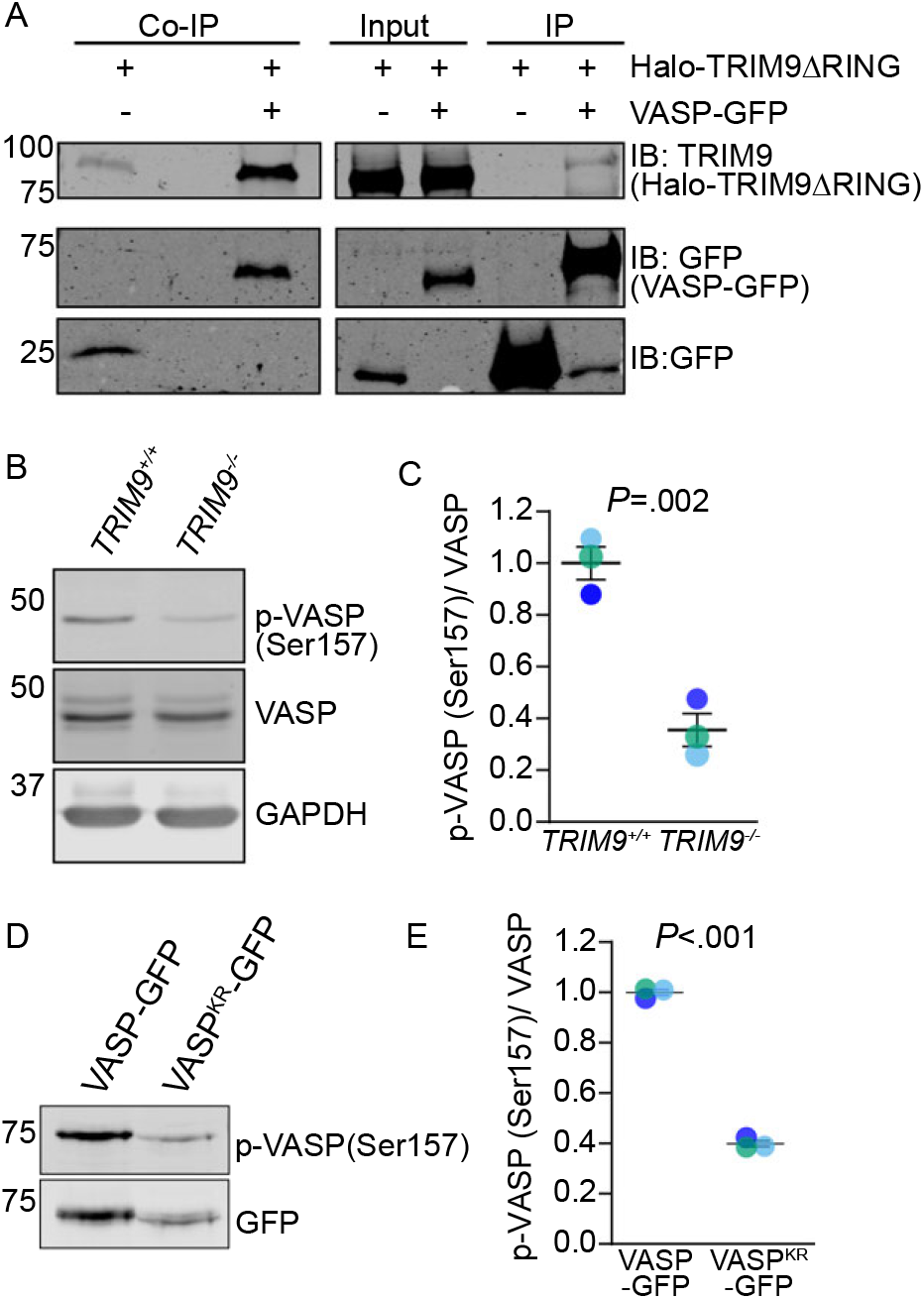
TRIM9 interacts with VASP in human melanoma and increases VASP phosphorylation. (**A**) Co-immunoprecipitation assays (Co-IP) from Dithiobis (succinimidyl propionate) DSP-crosslinked lysates from 1205Lu cells transfected with VASP-GFP or GFP and transduced with an adenoviral HALO-TRIM9ΔRING, n = 3 independent biological replicates. (**B**) Representative western blot in 1205Lu lysates of total VASP and phospho-Ser157 VASP in *TRIM9*^*+/+*^ cells. GAPDH is a loading control (**C**) Quantification of phospho-VASP (p-VASP Ser157) normalized to total VASP levels in 1205Lu cells; loss of TRIM9 reduced levels of phospho-VASP (*P* =.002, paired two-tailed *t*-test), n = 3 independent biological replicates. (**D**) Representative western blot of 1205Lu *TRIM9*^*+/+*^ cells transiently expressing VASP-GFP or VASP^KR^-GFP probed for GFP and phospho-VASP (Ser157). (**E**) Quantification of phospho-VASP (Ser157) normalized to total VASP-GFP or VASP^KR^-GFP revealed decreased p-VASP of VASP^KR^-GFP as compared to control VASP-GFP (*P* <.001, two-tailed paired *t*-test), n = 3 independent biological replicates.

Immunofluorescence suggested there was increased tritoninsoluble VASP protein in *TRIM9*^−/−^ cells (**Fig 3A-B**). However, analysis of VASP protein after stringent SDS lysis did not detect differences in VASP protein in *TRIM9*^*+/+*^ or *TRIM9*^−/−^ cells (**Fig 5B**,**Fig S5B**). This suggested in the absence of TRIM9, VASP exhibited increased association with the triton-insoluble cytoskeleton. Consistent with this, we previously demonstrated that ubiquitylation of VASP decreased F-actin binding (McCormick et al., 2024b). Similarly, VASP phosphorylation at Ser157 alters filament binding (Harbeck et al., 2000; Walders-Harbeck et al., 2002), enhancing localization of VASP to lamellipodia, presumably at barbed actin ends (Benz et al., 2009). Ser157 phosphorylation of VASP produces a mobility shift from ~ 46 kDA to ~ 50 kDA by SDS-PAGE (Wentworth et al., 2005; Döppler et al., 2013; Benz et al., 2009; Butt et al., 1994). Loss of TRIM9 reduced p-VASP (Ser157) levels (**Fig 5B-C**). We hypothesized that ubiquitylation of VASP may alter phosphorylation of Ser157. To test this hypothesis, we utilized a VASP^KR^ construct in which 9 lysine residues were mutated to arginine, resulting in reduced TRIM9-dependent VASP ubiquitylation (Menon et al., 2015). We expressed VASP-GFP or VASP^KR^-GFP in 1205Lu *TRIM9*^*+/+*^ cells and evaluated VASP phosphorylation at Ser157 (**Fig 5D**). This revealed reduced Ser157 phosphorylation on VASP^KR^ (**Fig 5E**), suggesting ubiquitylation preceded or promoted phosphorylation. Taken together these data suggested that TRIM9 interacted with and regulated VASP PTMs in melanoma.

### TRIM9 regulates VASP and zyxin dynamics at focal adhesions

To investigate how TRIM9 altered VASP behavior at focal adhesions, we employed fluorescence recovery after photobleaching (FRAP) of GFP-VASP in focal adhesions behind the leading edge of polarized cells (**Fig 6A**, Video S3). Fluorescence intensity was monitored until it reached a plateau, indicating steady state recovery (**Fig 6B**). Loss of TRIM9 did not affect the plateau of FRAP, indicating the mobile and immobile fractions of VASP were unaltered (**Fig 6B, Fig S6A**). However, loss of TRIM9 accelerated the recovery of GFPVASP fluorescence (**Fig 6C**), resulting in a smaller t_1/2_ (**Fig 6D**). This indicated that TRIM9 slowed VASP dynamics in focal adhesions. We questioned whether dynamics of other focal adhesion proteins were affected by TRIM9. VASP interacts with a subset of proteins in focal adhesions, including zyxin (Reinhard et al., 1995; Drees et al., 2000), but is not known to interact with paxillin. The rates of both zyxin and paxillin displayed behavior similar to VASP, with more rapid turnover (**Fig 6E-F, Fig S6B-C**). In contrast to VASP, paxillin and zyxin exhibited a small, but significantly increased amount of FRAP in *TRIM9*^−/−^ cells (**Fig S6D**).

**Fig. 6.**
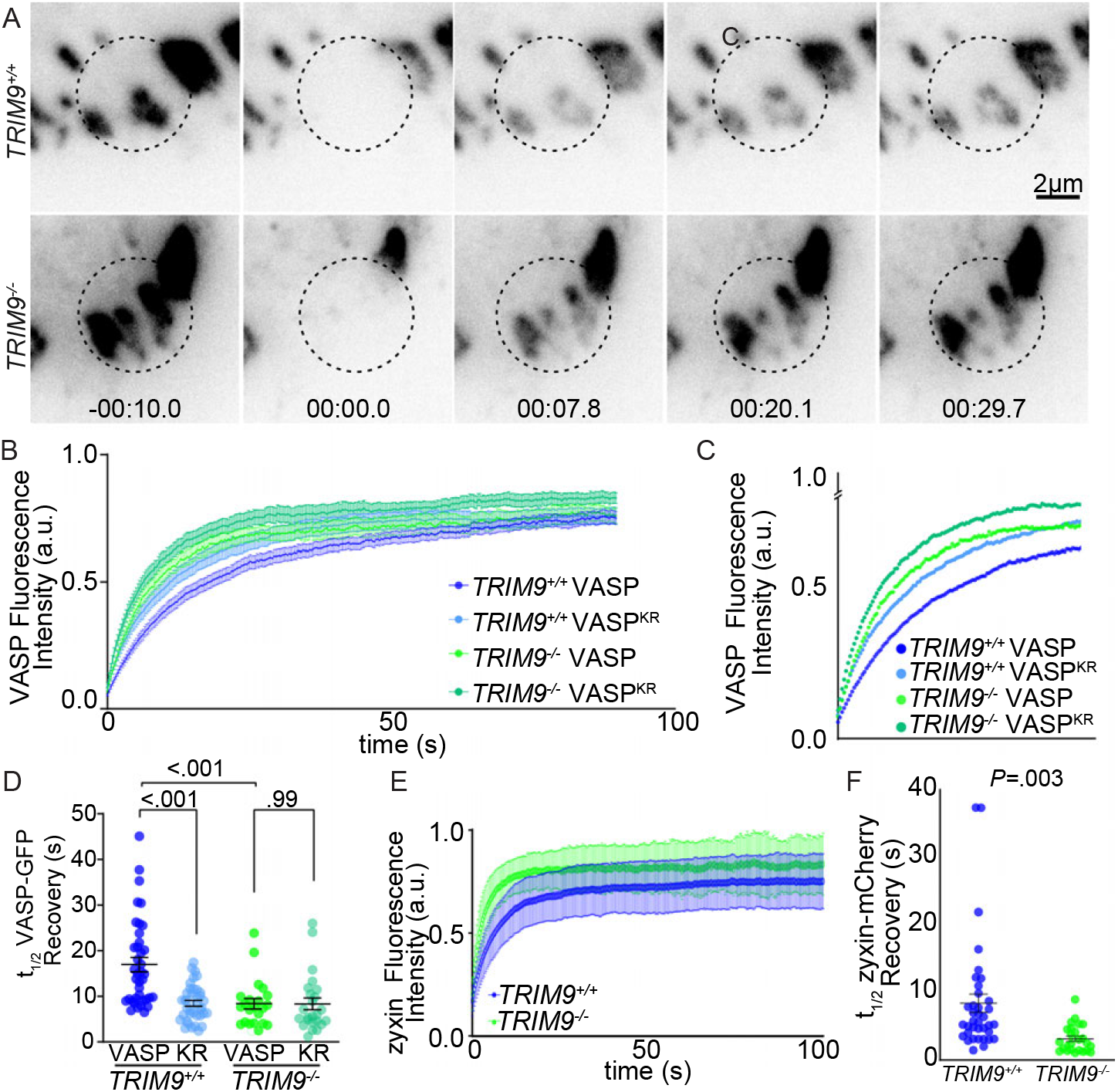
TRIM9 regulates VASP and Zyxin dynamics at focal adhesions. (**A**) Representative images of FRAP experiment of VASP-GFP at focal adhesions in *TRIM9*^*+/+*^ and *TRIM9*^−/−^ cells. (**B**) A normalized fluorescence intensity of VASP-GFP and VASP^KR^-GFP after photobleaching, n = 3 biological replicates; n 25 cells (**C**) Zoom of fluorescence intensity of VASP-GFP and VASP^KR^-GFP after photobleaching to emphasize differences in t_1/2_ of (B). (**D**) t_1/2_ of VASP-GFP and VASP^KR^-GFP in 1205Lu *TRIM9*^*+/+*^ and *TRIM9*^−/−^ cells; each dot represents the recovery rate of an individual cell. n = 3 independent biological replicates. *P*-values from One-way ANOVA (Bonferroni’s multiple comparisons test), n = 3 biological replicates; n ≥ 22 cells per condition. (**E**) A normalized fluorescence intensity of zyxin-mCherry indicated moderately increased zyxin fluorescence recovery due to loss of TRIM9 (S6D *P* =.01), Two-tailed *t*-test n = 3 biological replicates; n ≥ 25 cells. (**F**) t_1/2_ kinetics of zyxin-mCherry in 1205Lu cells (*P* =.0026, two-tailed *t*-test), n = 3 biological replicates; n ≥ 25 cells.

To evaluate whether ubiquitylation of VASP altered turnover dynamics, we employed VASP^KR^ in FRAP assays (**Fig 6B-D**). VASP^KR^ exhibited accelerated recovery dynamics in *TRIM9*^*+/+*^ cells (smaller t_1/2_), similar to GFP-VASP in *TRIM9*^−/−^ cells. This was consistent with the conclusion that TRIM9-dependent ubiquitylation of VASP slowed VASP turnover at adhesions. Supporting this conclusion, the fluorescence recovery of VASP^KR^ was indistinguishable from GFP-VASP in *TRIM9*^−/−^ cells. These data suggested that TRIM9 altered the dynamics of a subset of proteins in focal adhesions, perhaps through ubiquitylation of VASP. Focal adhesion Kinase (FAK) is a non-receptor tyrosine kinase activated early in focal adhesion formation that regulates focal adhesion formation, maturation, and disassembly. Although neurons do not exhibit large focal adhesions, we previously identified T RIM9 a s a r egulator o f FAK a ctivity (Plooster et al., 2017; Mutalik et al., 2025). To determine whether TRIM9 may also regulate adhesions via altered FAK signaling, we quantified levels of auto-phosphorylated and activated FAK (pY397) in *TRIM9*^*+/+*^ and *TRIM9*^−/−^ cells but found no substantial or significant alteration (**Fig S6E-F**). Together, these data suggested that TRIM9 modulated VASP localization and focal adhesion dynamics via direct regulation of VASP.

### Loss of TRIM9 alters cell morphology, collective cell contractility, and migration

We hypothesized that TRIM9-mediated alterations in focal adhesions and actin bundles altered the contractility and motility of melanoma cells. To evaluate collective cell contractility, we utilized a collagen gel assay, in which cells were embedded in and contract type-1 collagen. Whereas collagen gels without cells maintained their area, both *TRIM9*^*+/+*^ and *TRIM9*^−/−^ cells contracted collagen gels, reducing the area substantially after 24 hours (**Fig 7A**). However, *TRIM9*^−/−^ cells reduced the area significantly more. This result indicated that loss of TRIM9 increased cell contractility, corresponding with a more adhesive and bundled actin phenotype.

**Fig. 7.**
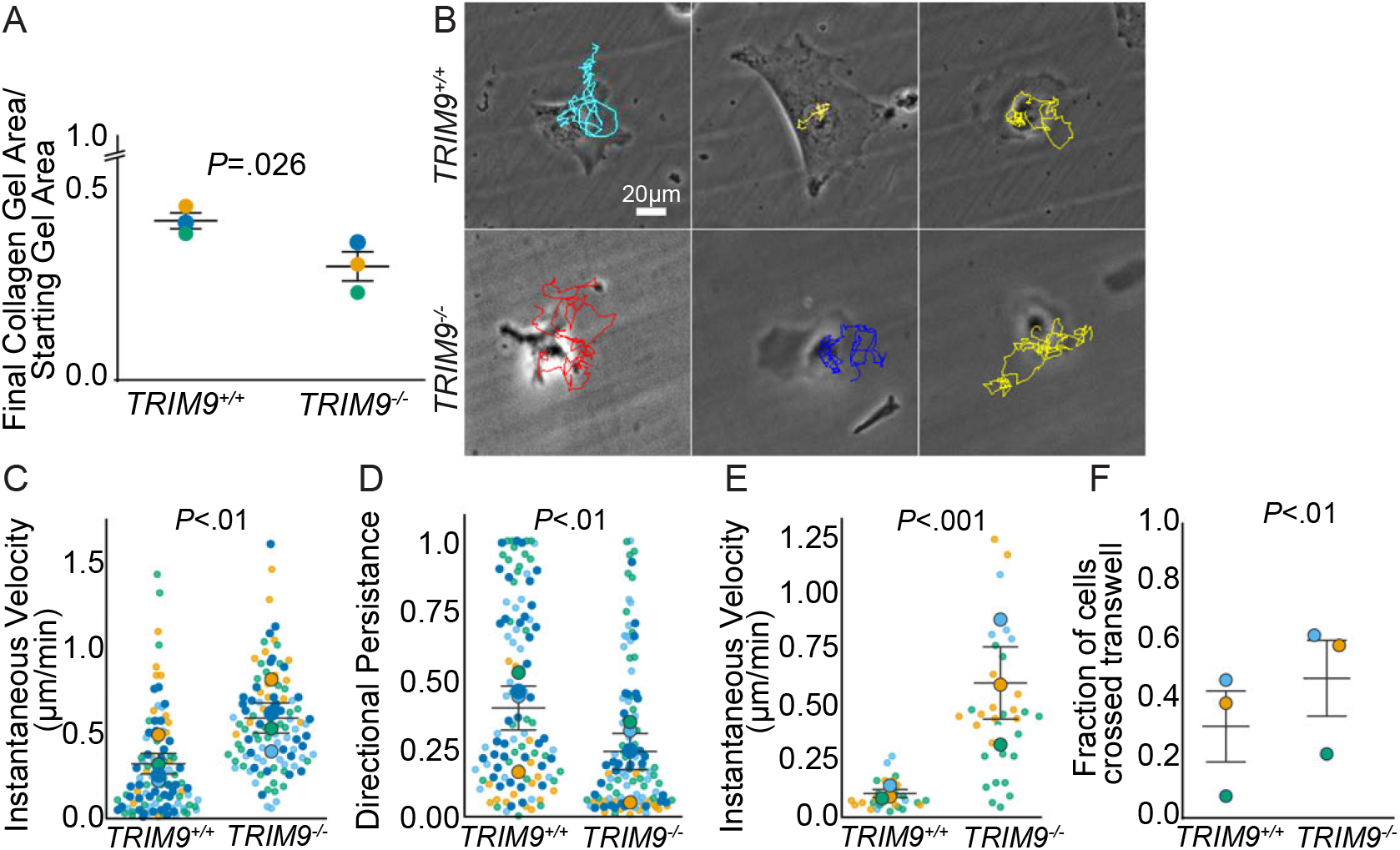
TRIM9 negatively regulates cell contractility, migration, and invasion in vitro. (**A**) 1205Lu cells were embedded in collagen gels for 24 hours and the difference in collagen gel area was quantified; TRIM9 loss increased the reduction of gel area (*P* =0.026, One way ANOVA with Dunnet’s multiple comparisons test), n = 3 independent biological replicates with a minimum of three technical replicates per experiment. (**B**) 1205Lu cells plated on laminin coated coverslips imaged for 18 hours using phase contrast time lapse imaging. Lines represent migration of cells over imaging duration. (**C**) The instantaneous velocity of *TRIM9*^*+/+*^ cells decreased compared to *TRIM9*^−/−^ cells plated on laminin (*P* <0.01, paired-two tailed *t*-test), n = 4 independent biological replicates, each color is a replicate; n ≥ 15 cells per genotype per replicate, each dot is a cell. (**D**) The directional persistence of 1205Lu *TRIM9*^−/−^ cells decreased compared to *TRIM9*^*+/+*^ cells (*P* <.01, paired-two tailed *t*-test), n = 4 independent biological replicates, each color is a replicate; n ≥ 15 cell per replicate, each dot is a cell. (**E**) Instantaneous velocity of 1205Lu cells plated on soft hydrogels; loss of TRIM9 increased instantaneous velocity (*P* <.001, paired-two tailed *t*-test), n = 3 independent biological replicates, each color is a replicate; n ≥ 30 cells, each dot is a cell. (**F**) 1205Lu cell Transwell cell invasion; loss of TRIM9 increased cellular invasion (*P* <.01, paired-two tailed *t*-test), n = 3 independent biological replicates, with a minimum of two technical replicates per experiment.

To determine the effects of loss of TRIM9 on cell motility, we performed random migration assays in 1205Lu cells plated on laminin (**Fig 7B**, Video S4). Loss of TRIM9 resulted in increased instantaneous cell velocity (**Fig 7C**) and decreased directional persistence of migration (**Fig 7D**). We found similar results in 1205Lu cells plated on fibronectin and PDL coated cover glass and uncoated cover glass (**Fig S7**). *TRIM9*^−/−^ cells consistently demonstrated an increased instantaneous velocity (**Fig S7A**) and decreased persistence (**Fig S7B**). We found that loss of TRIM9 also increased instantaneous velocity on soft hydrogels (**Fig 7E**, Video S1). Actomyosin contractility in cancer cells deforms and remodels the ECM in vivo, creating pro-invasive microenvironments that facilitate metastasis (Provenzano et al., 2008). We wondered whether directed chemotaxis through a matrix was altered by loss of TRIM9. We utilized transwell invasion assays incorporating basement membrane components to evaluate cell motility and remodeling of the ECM. Loss of TRIM9 increased the fraction of cells that invaded through the basement membrane (**Fig 7F**), suggesting that TRIM9 also negatively regulated directed chemotactic invasion. Taken together these data suggested TRIM9 was a novel regulator of melanoma cell contractility and motility.

### Loss of TRIM9 increases degradative capacity and reduces proliferation

Invasion involves proteolytic degradation of the ECM mediated by matrix metalloproteinases (MMPs). To investigate if the increased invasion of *Trim9*^−/−^ cells (**Fig 7F**) was coincident with increased degradative capacity, we performed gelatin degradation zymography assays using conditioned media from 1205Lu cells. Loss of TRIM9 increased the capacity of conditioned media to degrade gelatin (**Fig 8A-B**), suggesting high TRIM9 levels decreased secretion of MMPs. Immunoblotting indicated MMP2 levels increased 2.1-fold in conditioned media of *TRIM9*^−/−^ cells (**Fig 8C**). As netrin secretion has been implicated in multiple cancers including melanoma (Kaufmann et al., 2009; Boussouar et al., 2020), we also probed conditioned media for netrin-1 (**Fig 8D**). This revealed loss of TRIM9 increased netrin secretion by ~ 4.1 fold. Taken together, these data suggested TRIM9 was a negative regulator of secretion in melanoma. This aligns with increased exocytosis in developing neurons lacking TRIM9 (Winkle et al., 2014; Urbina et al., 2018).

**Fig. 8.**
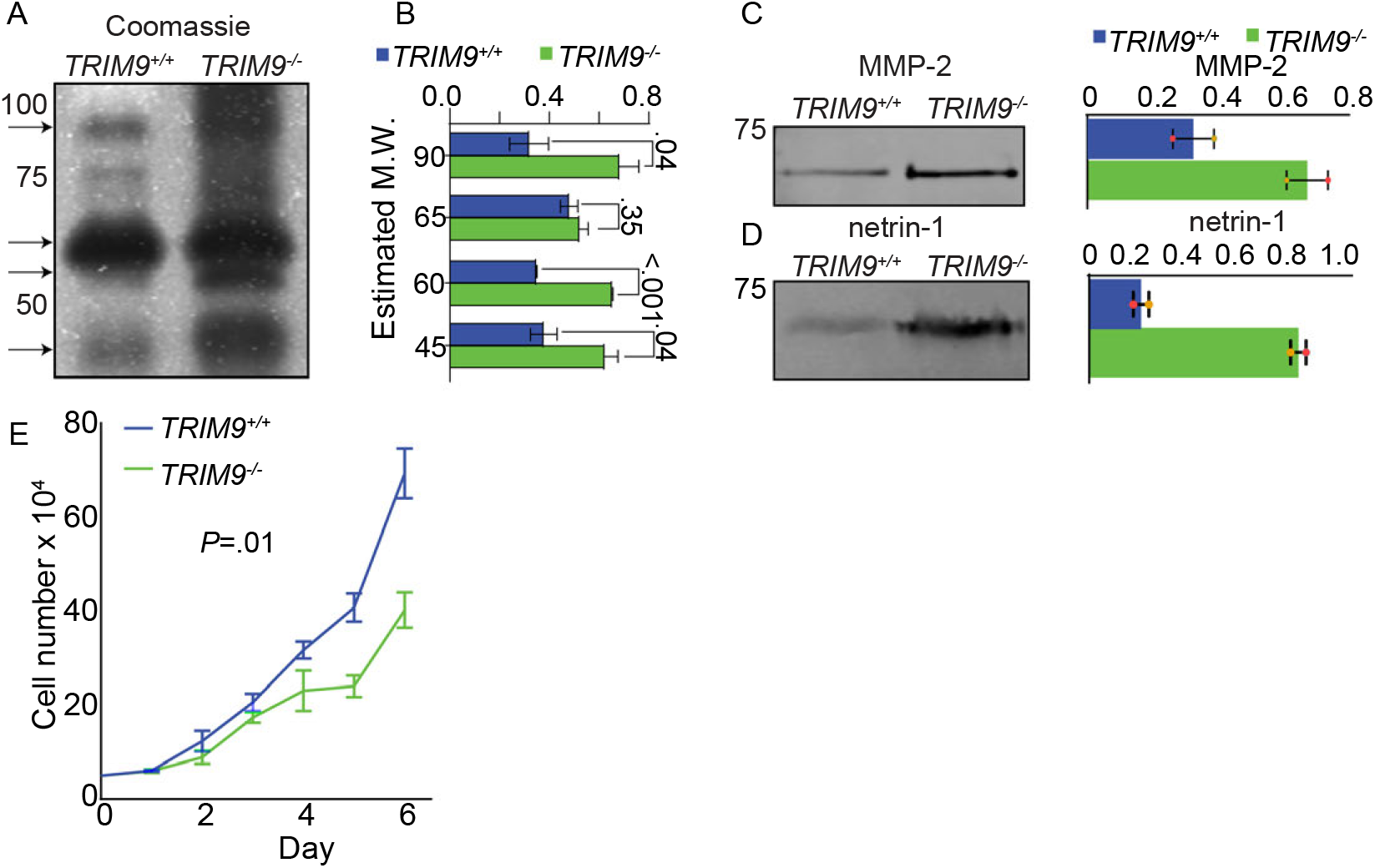
Loss of TRIM9 increases degradative secretion and reduces proliferation. (**A**) Zymography gel of conditioned media from serum starved 1205Lu cells. Loading controlled by cell count at time of conditioned media collection. (**B**) Quantification of bands at 90, 65, 60, 45 kDa (black arrows) indicated that loss of TRIM9 increased gelatin degradation (*P* =.04, <.001, .04, two-tailed *t*-test), n= 2 independent biological replicates. (**C**) Western blot and quantification of 1205Lu conditioned media immunoblotted for MMP-2. (**D**) Western blot and quantification of 1205Lu conditioned media immunoblotted for netrin-1. (**E**) Cell number counts over a 6-day time course; loss of TRIM9 reduced cell proliferation (growth rate constant k= *TRIM9*^*+/+*^ 0.437 ± 0.021 vs *TRIM9*^−/−^ 0.346 ± 0.028, *P* =.01, two-tailed *t*-test), n =3 independent biological replicates with 2 technical replicates per experiment.

Uncontrolled cell proliferation is a defining hallmark of cancer, and dysregulated growth underlies tumor progression and therapeutic resistance. Cell proliferation counts with exponential growth modeling revealed that *TRIM9*^*+/+*^ cells proliferated ~ 26% faster than *TRIM9*^−/−^ cells (**Fig 8E**). MTT based proliferation assays corroborated the increased proliferation of *TRIM9*^*+/+*^ cells (**Fig S8**). Notably, there were no differences observed in cell death in counting assays. These data suggested that TRIM9 promoted cell proliferation.

### TRIM9 promotes melanoma proliferation and alters metastasis in vivo

Our findings in 1205Lu and WM266-4 cells indicated TRIM9 was a potent regulator of melanoma phenotypes. High TRIM9 was associated with malignant properties such as increased blebbing and proliferation, whereas TRIM9 loss was associated with increased adhesion, increased contractility, increased secretion, and faster rates of mesenchymal migration. To evaluate TRIM9 in tumor progression in vivo, we turned to a model where we could induce an endogenous primary melanoma on the ear of a mouse with an intact immune system. The *Tyr:Cre*^*ER*^: *Pten*^*f/f*^*:Braf*^*CA*^*:tdTomato*^*LSL*^ (*PBT)* model harbors two common melanoma drivers: a floxed allele for the tumor suppressor PTEN (*P)* and a Cre-driven constitutively active BRAF (V600E) mutation (*B*), as well as a flox-stop-flox tdTomato reporter (*T*) and a tamoxifen-inducible Cre under a melanocyte specific Tyrosinase promoter (*Tyr:Cre*^*ER*^) (Brighton et al., 2018; Tagliatela et al., 2020; Dankort et al., 2009; Madisen et al., 2010). Previous RNAseq revealed that *Trim9* mRNA is expressed in the tumors in this model (Brighton et al., 2018). We validated enrichment of TRIM9 protein by immunoprecipitation and mass spectrometry. We crossed the *PBT* mice with *Trim9* floxed mice (*Trim9*^*f/f*^, **Fig 9A, Fig S9A**)(Winkle et al., 2014) and induced primary melanoma tumors lacking TRIM9 (*PBT-Trim9*^*f/f*^) and containing TRIM9 (*PBT-Trim9*^*+/+*^) with a brief application of 4-hydroxytamoxifen (4-HT) on the ear. We tracked TdTomato expression (recombination) and tumor progression via fluorescence and transmitted light microscopy approaches (**Fig 9B**).

**Fig. 9.**
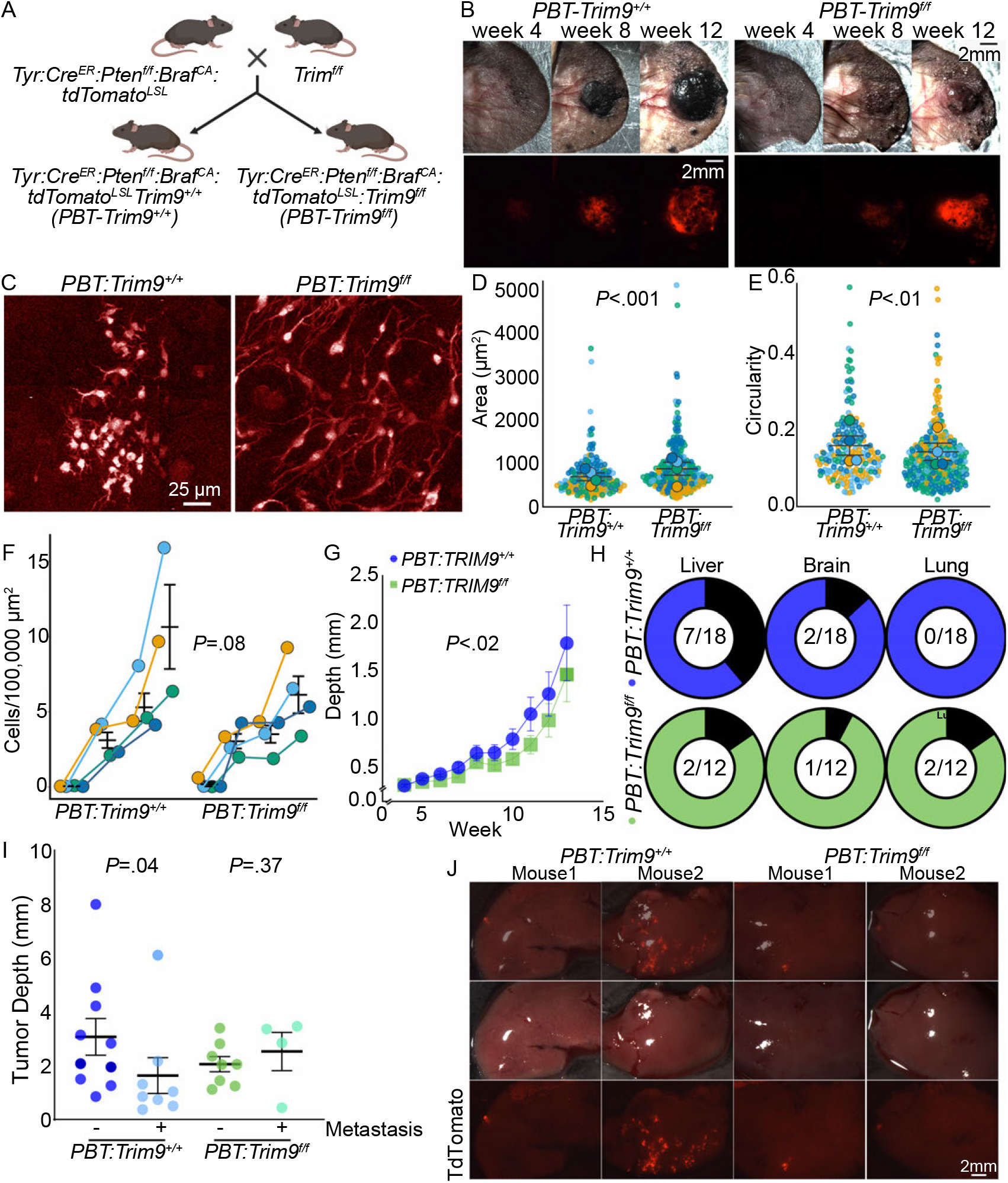
TRIM9 promotes tumor growth and alters metastasis in vivo. (**A**) Schematic of GEM models created by crossing a GEM harboring common melanoma drivers (*Pten*^*f/f*^ and a constitutively active *Braf*^*CA*^) with *TdTomato*^*LSL*^ to *Trim9*^*f/f*^ mice to generate *PBT-Trim9*^*+/+*^ and *PBT-Trim9*^*f/f*^. (**B**) Transillumination and fluorescence imaging (TdTomato) of the ear of *PBT-Trim9*^*+/+*^ and *PBT-Trim9*^*f/f*^ mice over a 12-week period following tumor induction. (**C**) Multi-photon image of TdTomato+ recombined cells in the primary tumor of *PBT-Trim9*^*+/+*^ and *PBT-Trim9*^*f/f*^ mice at five days post induction. Quantification of (**D**) cell area and (**E**) cell circularity from multi-photon imaging at day five post induction; TRIM9 loss increased cell area (*P* <.001, Mann-Whitney) and decreased cell circularity (*P* <.01, Mann-Whitney test), n = 4 mice per genotype, each color is a mouse. (**F**) Quantification of TdTomato+ cell density in the induction site one, five, eight, thirteen and fifteen days post induction; *Trim9*^*+/+*^ melanoma cells proliferated faster between day five and eight day (slope = 0.73) than *Trim9*^*f/f*^ melanoma cells (slope = 0.16; *P* =.08, Welch Two Sample *t*-test), n = 4 mice per genotype, each color is a mouse. (**G**) Exponential growth modeling of the mean thickness of the ear and primary tumor revealed tumor growth kinetics were reduced due to loss of TRIM9 (growth rates: *PBT-Trim9*^*+/+*^ 0.1381 mm/week, *PBT-Trim9*^*f/f*^ 0.0991 mm/week; *P* =.02, extra sum of squares F test, F (2,287) =3.744). n= 18 *PBT-Trim9*^*+/+*^, 13 *PBT-Trim9*^*f/f*^ mice. (**H**) Fraction of mice with TdTomato+ macro-metastases for each organ. (**I**) Tumor depth in mice with and without metastasis; there was an inverse relationship between tumor depth and metastases in *PBT-Trim9*^*+/+*^ mice (*P* =.04, Mann Whitney test). This relationship was not present in *PBT-Trim9*^*f/f*^ mice (*P* =.37, Mann Whitney test). (**J**) Example images of macro-metastases in the liver of two *PBT-Trim9*^*+/+*^ and two *PBT-Trim9*^*f/f*^ mice.

We first asked whether loss of TRIM9 in vivo affected tumor cell morphology. At 5 days post melanoma induction, TdTomato+ cells in the ear of *PBT-Trim9*^*f/f*^ mice were larger and less circular than in *PBT-Trim9*^*+/+*^ mice (**Fig 9C-E**), similar to findings in vitro (**Fig 1, Fig S1**). We next evaluated how TRIM9 loss affected cell proliferation. Functional data analysis revealed that *PBT-Trim9*^*+/+*^ cells exhibited a steady growth rate throughout the observation period (days 1–15), whereas *PBT-Trim9*^*f/f*^ cells displayed a biphasic growth pattern characterized by an early proliferative burst and a mid-period deceleration. A permutation test on the mean squared difference between smoothed growth curves approached statistical significance (*P* = 0.09, n = 4 per group). *PBT-Trim9*^*+/+*^ and *PBT-Trim9*^*f/f*^ tumors accumulated similar density of TdTomato+ cells at the induction site five days after recombination. To further characterize the period of early divergence, the slope of cell density change between days 5 and 8 was compared between groups. *PBT-Trim9*^*+/+*^ cells grew steadily (slope 0.73), whereas *PBT-Trim9*^*f/f*^ growth decelerated (slope 0.16, *P* =0.084, **Fig 9C-F**), similar to our in vitro findings (**Fig 8**,**Fig S8**). These data indicated TRIM9 also altered melanoma proliferation and morphology in vivo.

We next quantified tumor depth from 4 weeks to humane endpoints between 13 - 16 weeks post induction. Exponential growth modeling revealed that loss of TRIM9 was associated with a reduction in the rate of tumor growth (*PBT-Trim9*^*f/f*^: 0.091/mm/week; *PBT-Trim9*^*+/+*^: 0.1381mm/week) (**Fig 9G**), with primary tumors at 13 weeks significantly smaller in *PBT-Trim9*^*f/f*^ mice (*P*=.02, extra sum of squares F-test). We noted high variability in the size and appearance of both *PBT-Trim9*^*f/f*^ and *PBT-Trim9*^*+/+*^ primary tumors (**Fig S9B, Video S5-6**). Within-group variability increased substantially over time in both genotypes. In *PBT-Trim9*^*+/+*^ tumors, the coefficient of variation (CV) rose from 20.1% at week 4 to 91.9% at week 13, indicating marked divergence in individual tumor growth trajectories over time. *PBT-Trim9*^*f/f*^ tumors showed a similar but attenuated pattern, with CV increasing from 18.7% at week 4 to 68.9% at week 13. The greater trajectory divergence in *PBT-Trim9*^*+/+*^ tumors at late timepoints suggested that TRIM9 loss constrained the heterogeneity of tumor growth dynamics.

The brain, lungs, and liver, are clinically relevant areas of melanoma metastasis (Damsky et al., 2011; Conway et al., 2022). Macroscopic examination of TdTomato+ signal on the surface of these organs was performed to evaluate metastasis (**Fig 9H,J, Fig S9C**). Livers showed the most frequent metastasis (38.9% *PBT-Trim9*^*+/+*^, 16.7% *PBT-Trim9*^*f/f*^), followed by brain (11.1% *PBT-Trim9*^*+/+*^, 8.3% *PBT-Trim9*^*f/f*^), and lung (0% *PBT-Trim9*^*+/+*^, 16.7% *PBT-Trim9*^*f/f*^). In *PBT-Trim9*^*+/+*^ tumors, there was an inverse correlation between tumor depth and metastasis; mice with smaller primary tumors displayed increased metastasis to internal organs (**Fig 9I**). There was no apparent relationship between tumor depth and metastasis to internal organs in *PBT-Trim9*^*f/f*^ mice (**Fig 9I**). We noted varying degrees in the size and number of Td-Tomato+ foci between *PBT-Trim9*^*+/+*^ and *PBT-Trim9*^*f/f*^ mice; broadly *PBT-Trim9*^*+/+*^ mice tended to have larger metastases or more metastatic foci in comparison to *PBT-Trim9*^*f/f*^ mice (**Fig 9J, Fig S9C**). Notably, local metastasis on the ear occurred frequently in both *PBT-Trim9*^*f/f*^ mice (100%) and *PBT-Trim9*^*+/+*^ mice (78%). Taken together, these data suggested that TRIM9 promoted melanoma proliferation and influenced the dissemination, destination, and growth of metastasis.

## Discussion

Melanoma is a malignant and heterogenous cutaneous cancer with distinct cellular phenotypes associated with differential gene expression (Pozniak et al., 2024; Wouters et al., 2020; Andrews et al., 2022). These studies found that TRIM9 expression was high in the intermediate or neural crest-like states (Pozniak et al., 2024; Wouters et al., 2020; Andrews et al., 2022). Another study suggested TRIM9 expression is a potential prognostic marker in melanoma (Kutt et al., 2021). Although high TRIM9 expression correlates with poor patient survival (Kovács et al., 2023), how TRIM9 altered melanoma cellular phenotype was unclear. Our data suggest that the increased TRIM9 expression observed in melanoma alters cell proliferation, morphology, and motility. The inverse correlation between growth and metastasis of *PBT-Trim9*^*+/+*^ tumors, which was absent in *PBT-Trim9*^*f/f*^ tumors, may suggest that TRIM9 expression can promote phenotype switching between proliferative and migratory states. We suggest that TRIM9 is thus poised to alter melanoma phenotypes and disease progression.

### TRIM9 switches melanoma cells from a mesenchymal adhesion-based morphology to bleb-based morphology

We found that TRIM9 is a novel and potent regulator of melanoma cell morphology in vitro and in vivo, with TRIM9 loss resulting in a more polarized and spread cell state. Both 1205Lu and WM266-4 TRIM9 expressing cells exhibited circular, bleb-like morphology and reduced cellular area. This TRIM9-dependent switch between blebbing and mesenchymal states likely signifies changes in actin organization and adhesion or changes in hydrostatic cytoplasmic pressure.

Blebbing cells exhibit high cortical actomyosin contractility and low adhesion (Paluch et al., 2006; Tinevez et al., 2009). In contrast, mesenchymal migration requires tightly coupled dynamics of adhesions, F-actin based protrusions, and actomyosin contractility (Gardel et al., 2010; Gupton and Waterman-Storer, 2006). We found that TRIM9 loss enhanced cell adhesion, altered actin organization and promoted contractility, consistent with reduced blebbing. Alternatively recent work suggested soluble Ena/VASP proteins promote blebbing (Fujii et al., 2026) by altering osmotic pressure in blebs (Fujii et al., 2026). Our finding that TRIM9 loss promotes VASP integration into the cytoskeleton, effectively reducing soluble VASP, is also consistent with this possibility. Further investigations are needed to understand how high TRIM9 levels promote blebbing.

TRIM9 loss increased adhesion, contractility, and migration as well as degradative capacity and chemotactic invasion in vitro. These phenotypes might predict loss of TRIM9 to increase metastasis, however the effect of TRIM9 loss on metastases was complex. We observed increased local metastases to the ear and macro-metastases to the lung in the absence of TRIM9, but decreased and smaller metastases to liver and brain. In melanoma, the transition from mesenchymal to bleb-based amoeboid states, can increase the speed of motility in confined spaces (Taddei et al., 2014; Gabbireddy et al., 2021; Behrooz and Shojaei, 2024). Several studies indicate bleb-based motility of melanoma is proteolytically independent (Driscoll et al., 2024; Gabbireddy et al., 2021). We found that high TRIM9 decreased secretion in vitro and promoted a bleb like morphology in vitro, and a smaller, less polarized morphology in vivo. More studies are needed to determine how TRIM9 alters melanoma cell morphology and migratory modes in vivo, as well as matrix remodeling and proteolytic activity, and whether this influences the frequency and sites of metastasis and size of metastatic foci.

In addition to altered methods of migration and invasion, promotion of bleb-like morphology in melanoma prevents anoikis, cell death due to substrate detachment, via formation of oncogenic signaling hubs (Weems et al., 2023). Preventing anoikis may confer survival advantage in vivo after intravasation, while cancer cells are in the circulatory system. TRIM9 positive melanoma cells proliferated more and displayed more frequent organ metastasis, which suggests cell signaling and interactions with the immune system may be altered. Whether TRIM9 mediated bleb promotion results in altered cell signaling, survival, metastatic seeding, or immune evasion in vivo will be important to consider in future studies. Of note, high TRIM9 expression in patient tumors correlated with poor overall survival and a reduced response to anti-PD-1 and anti-CTLA-4 immunotherapy.

### TRIM9 alters VASP localization, modification, and dynamics at focal adhesions

This function of TRIM9 in melanoma differs from the role of TRIM9 in the growth cone of developing neurons. In neurons, TRIM9 loss increased filopodia (Menon et al., 2015).

However, neurons lack prominent focal adhesions and stress fibers and rather form small focal contacts. These differences in actin architecture and adhesion may underlie key differences in cell type specific effects of TRIM9 on adhesion and actin organization. Lamellipodia, highly protrusive Arp2/3 driven branched F-actin structures involved in mesenchymal migration (Nicholson-Dykstra and Higgs, 2008; Wu et al., 2012; Nolen et al., 2009) were thinner upon TRIM9 loss. VASP also localizes to the edge of lamellipodium and in mouse B16-F1 melanoma cells, VASP depletion similarly reduced lamellipodial width (Damiano-Guercio et al., 2020). This may suggest that in *TRIM9*^−/−^ melanoma cells the preferential localization of VASP to focal adhesions has an adverse effect on lamellipodia morphology. However, absolute VASP levels in lamellipodia increased, which may compete with Arp2/3 and alter filopodia (Skruber et al., 2020; Damiano-Guercio et al., 2020; Tang et al., 2025). In developing neurons, loss of TRIM9 reduced Arp2/3 levels in the postsynaptic density (McCormick et al., 2024a). However, whether TRIM9 intersects with Arp2/3 function independent of VASP in various cell types is unclear.

Our findings suggest that VASP alters both actin architecture and focal adhesion phenotypes in melanoma. Loss of TRIM9 increased VASP incorporation into triton-insoluble cytoskeletal structures. This is consistent with our previous work demonstrating TRIM9 multi-monoubiquitylates VASP in a nondegradative manner to alter VASP localization and reduce the ability of VASP to bind, bundle, and accelerate the elongation of actin filaments (Menon et al., 2015; Boyer et al., 2020; McCormick et al., 2024b). Phosphorylation of VASP Ser157 also alters VASP actin binding and promotes VASP localization to the leading edge of spreading cells (Döppler et al., 2013; Wentworth et al., 2005; Benz et al., 2009). Loss of TRIM9 and VASP^KR^ expression reduced Ser157 phosphorylation. Together, these findings suggested that TRIM9-mediated ubiquitylation may promote Ser157 phosphorylation. VASP ubiquitylation may influence structural changes that enhance access of kinases, such as PKA, PKC or PKD, to Serine157. Alternatively, VASP ubiquitylation may regulate dwell time or localization at subcellular structures where these kinases function, for example at focal adhesions, as suggested by our FRAP studies. PKA, PKC, and PKD regulate focal adhesion dynamics (Kang et al., 2024; McKenzie et al., 2020; di Blasio et al., 2015). How the combination of PTMs alter VASP localization and activity at focal adhesions is an intriguing question.

Loss of TRIM9 and altered VASP localization enhanced stress fiber bundling and increased focal adhesion number and elongation, characteristics associated with contractility and commonly found in migratory mesenchymal cells (Buskermolen et al., 2018; Desjardins-Lecavalier et al., 2023). Loss of TRIM9 accelerated the turnover of VASP, zyxin, and paxillin within adhesions. Zyxin is a LIM domain containing protein that localizes to stress fiber strain sites, where it recruits VASP and α actinin to promote polymerization and crosslinking (Smith et al., 2010; Phua et al., 2025). Zyxin recruits VASP to focal adhesions in a tension-dependent manner, which promotes actin polymerization (Sala et al., 2025). We anticipate in the absence of TRIM9, stress fibers were likely under increased strain, which may enhance recruitment of VASP and α actinin and increase actin polymerization and bundling. Alternatively or in addition, altered focal adhesion composition may promote contractility. In endothelial and airway smooth muscle cells, VASP overexpression increased actomyosin contractility through its interaction with vinculin (Furman et al., 2007; Wu and Gunst, 2015). In contrast, loss of VASP and zyxin reduced focal adhesion number, area, and the density, altered actin polarity and adhesions, and reduced the number of dorsal and radial fibers in mouse embryonic fibroblasts (Chung et al., 2025). TRIM9 may regulate the complex interplay of VASP functions at focal adhesions and actin protrusion to both alter contractility and actin polymerization in non-neuronal motile cells is an intriguing possibility. Loss of TRIM9 function in zebrafish macrophages reduced motility (Tokarz et al., 2017), but whether this was dependent on altered VASP function is unknown. In highly invasive breast cancer cells, elevated VASP expression promotes migration and invasion (Tian et al., 2018). Further, expression of VASP correlates with poor prognosis in gastrointestinal cancer patients and promotes liver metastasis (Xiang et al., 2018). As TRIM9 is found in many cancer types, investigating TRIM9-dependent regulation of VASP may be warranted.

### Netrin-TRIM9 signaling axis in melanoma

Deletion of TRIM9 increased secretion of netrin-1 from melanoma cells in vitro, but how this may alter melanoma phenotypes and the tumor microenvironment is unknown. From our work in neurons, we know that TRIM9 interacts with the DCC and UNC5 (Winkle et al., 2014; Mutalik et al., 2025), receptors for the axon guidance cue netrin-1, and that TRIM9 regulates netrin-dependent neuronal morphogenesis. Upregulation of netrin-1 and loss of the netrin receptor DCC are protumoral in many cancers, including melanoma (Rodrigues et al., 2007; Kaufmann et al., 2009; Mehlen and Guenebeaud, 2010; Mehlen et al., 2011; Castets et al., 2011; Krauthammer et al., 2012; Boussouar et al., 2019). Upregulation of the repulsive UNC5 netrin receptors is also implicated in tumor progression (Mehlen and Guenebeaud, 2010; Castets et al., 2011; Zhou et al., 2022). In the context of melanoma, netrin can promote invasion, migration, and progression (Kaufmann et al., 2009; Boussouar et al., 2020). In murine models of melanoma, netrin is secreted by more metastatic tumors (Boussouar et al., 2020). NP137 is a netrin function blocking antibody that prevents epithelial to mesenchymal transition and reduces migration in carcinoma cells and murine tumor models (Lengrand et al., 2023; Cassier et al., 2023). NP137 is currently in clinical trials for a variety of solid cancers, and clinical trials are evaluating synergism with anti-PD1/PDL-1 in patients who have previously had poor response to these immunotherapies (Centre Leon Berard, 2026). Although it remains unclear whether TRIM9 functions downstream of netrin-1 in melanoma, our findings raise the possibility that netrin-dependent modulation of TRIM9 activity regulates melanoma phenotypes and the switch between bleb-based to adhesive states resulting in increased metastasis in vivo.

## Materials and Methods

### Cell Culture

1205Lu and WM266-4 cells were cultured in DMEM (Gibco 11965092) with 10% FBS (Fisher FB12999102) and maintained at 5% CO_2_, 37°C. For passaging and plating, cells were washed twice with DPBS followed by incubation with 0.25% trypsin EDTA (Gibco 25200072) at 37°C. An equal volume of culture media was added to trypsin-EDTA to quench trypsin reaction and cells were plated at desired density. WM266-4 cells were a gift from Ned Sharpless (UNC Chapel Hill). 1205Lu cells were a gift from Simon Rothenfusser (Munich); identification was verified via STR profiling by LabCorp. Cells lines were routinely tested for mycoplasma by qPCR through the UNC Vironomics Core.

### Knockout Generation

1205Lu *TRIM9*^−/−^ cells were generated by CRISPR/Cas9 gene editing using a guide RNA (gRNA) targeting the sequence 5’-GCG GCT ATG GCT CCT ACG GGG GG-3’ in exon 1 of the human *TRIM9* gene. 5 x 10^5^ cells were transfected in 6-well plates with the expression plasmid (3 µg DNA/9 µl TranIT transfection reagent (Mirus Bio, Madison, WI)) coding for a CMV driven CAS9-2A-RFP cassette and the U6 promoter driven guide RNA. 24 hours after transfection, RFP-positive cells were sorted cloned by limiting dilution and expanded. Genomic DNA was extracted using QuickExtractTM DNA extraction solution (Epicentre) and the target region of interest was amplified by PCR (forward primer 5’ CACAGAGCTAGCGCCTCTC-3’; reverse primer 5’-TACGCGATTCTTGGGGAAGC-3’). After prescreening, using the mismatch-sensitive T7 endonuclease I (T7EI), positive clones in the T7 assay were sequenced. Knockout-cell clones were identified as cell clones harboring biallelic frameshift mutations. Genotypes of the respective knockout cell lines are available upon request. Lack of TRIM9 expression was confirmed by western blot.

WM266-4 *TRIM9*^−/−^ cells were generated by CRISPR/Cas9 lentiCRISPR V2 (Addgene #52961) using gRNA targeting the sequence 5’-GCG GCT ATG GCT CCT ACG GGG GG-3’ in exon 1 of the human *TRIM9* gene. Lentivirus was produced in HEK293FT cells using packaging plasmids pRSV-rev (Addgene 12253), pMDLg/pRRE-(Addgene#12251), pCMV-VSV-G (Addgene#8454). Media containing virus was filtered using 0.2 µm filter and subsequently cells were transduced with 4 µg/mL polybrene (Sigma TR1003). 48 h post transduction cells were grown in 2.5 µg/ml puromycin (Thermo J67236XF). Post puromycin selection, 1 µg/ml of doxycycline (Sigma D9891) was added to induce GFP-Cas9 activation. Cells were sorted using flow cytometry (Biorad S3 Cell Sorter) for GFP-Cas9 positive cells and subsequently sorted until only a pure GFP-Cas9 positive population was achieved. Cells were kept under doxycycline induction and puromycin selection until a pure TRIM9 knockout population was achieved. Cells were subsequently immunoblotted for TRIM9 using an inhouse generated antibody (Winkle et al., 2014) and anti TRIM9 (Proteintech 10786-1-AP rabbit) antibody. Genetic knockout was confirmed by sequencing (Azenta), with complex indels inserted causing a frameshift insertion.

### Mouse

The *Tyr:Cre*^*ER*^: *Pten*^*f/f*^*:Braf*^*CA*^*:tdTomato*^*LSL*^ (*PBT)* model was generated previously (Brighton et al., 2018; Tagliatela et al., 2020; Dankort et al., 2009; Madisen et al., 2010). In these mice, a tamoxifen inducible CreERis expressed under the melanocyte-specific promoter Tyrosinase (*Tyr:Cre*^*ER*^). Endogenous *Pten* is flanked by LoxP sites on both alleles, endogenous WT *Braf* cDNA is flanked by LoxP sites and followed by *Braf* V600E mutant expressed from the endogenous promoter upon deletion of the WT cDNA. A LoxP/stop/LoxP site precedes TdTomato, which upon recombination induces TdTomato expression. The *PBT-Trim9*^*f/f*^ model was derived from crossing *PBT* model with *Trim9*^*f/f*^ mice (Menon et al., 2015). Breeding, care, and maintenance of each strain were carried out in accordance to the UNC Division of Comparative Medicine and Institutional Animal Care and Use Committee-approved protocols and standards. Genotyping was completed from DNA isolated from ear clips of pups before use in experimentation in house by PCR and via Transnetyx.

Tumors were induced in *PBT-Trim9*^*+/+*^ and *PBT-Trim9*^*f/f*^ mice by local application of 4-HT (Sigma; H6278) to the right ear of 12-14 week-old mice of both sexes as described previously (Brighton et al., 2018; Tagliatela et al., 2020). Briefly, 1 µL of 20 mM 4-HT solution in DMSO, 100% ethanol, and orange-6 dye was applied under anesthesia for precisely ten minutes before the area was thoroughly washed with 70% ethanol for 2 minutes. Tumor growth was quantified by measuring the thickness of the ear where the tumor was induced every seven days by caliper while the mice were anesthetized with 1.5% isoflurane. Multiphoton images were acquired 1 day prior to induction and 1,5,8 days post induction while the mice were anesthetized with 1.5% isoflurane.

### Plasmids, Antibodies, Virus

1205Lu and WM266-4 cells were transfected with Lipofectamine 2000 (Thermo 11668019) according to the manufacturer’s instructions.

Plasmids: Lentivirus packaging plasmids used were pRSV-rev (Addgene 12253), pMDLg/pRRE-(Addgene#12251), pCMV-VSV-G (Addgene#8454). peGFP-N1-VASP (human) and peGFP-N1-VASP^KR^ were previously described and generated, in which K240,252,283,286,321,348,357,358,363 were all mutated to R (Menon et al., 2015). The HALO-TRIM9 construct was previously described and generated using pcs2-Myc-TRIM9 plasmid(Winkle et al., 2014; Mutalik et al., 2025). GFP-paxillin was acquired from James Bear. Zyxin m-Cherry was a generous gift from Irina Kavirina, Vanderbilt University.

Antibodies used : rabbit polyclonal anti GFP (A11122, Invitrogen, RRID:AB_221569), rabbit polyclonal anti TRIM9 (generated in house), rabbit polyclonal anti TRIM9 (Proteintech 10786-1-AP), chicken polyclonal anti-GFP (GFP-1010, Aves), mouse monoclonal anti paxillin (BD Transduction Laboratories 610051), mouse monoclonal anti GAPDH WB(Santa Cruz mouse (D-6): sc-166545), rabbit polyclonal anti VASP (Sigma-Aldrich rabbit Anti-VASP (C-terminal) antibody: V3390), rabbit anti VASP (Cell Signaling Technology antirabbit Phospho-VASP (Ser157) Antibody #3111), rabbit monoclonal anti phospho-FAK (Tyr397 (Invitrogen 44-625G), mouse monoclonal anti FAK (Invitrogen 39-6500), Alexa Flour Phalloidin 488 (Fisher A12379), Alexa Flour Phalloidin 568 (A12380). AF anti mouse 647 (A31571), AF anti rabbit 568 (A10042). Licor goat anti rabbit 680 (Licor 926-32211), Licor goat anti mouse 680 (Licor 926-32210).

### Cell Lysis, Co-Immunoprecipitation, and Western Blotting

Cells were lysed in 1× RIPA buffer (50 mM Tris-HCl (pH 7.4), 150 mM NaCl, 1% NP-40, 0.5% sodium deoxycholate, 0.1% sodium dodecyl sulfate (SDS), 1 mM EDTA, 15 mM sodium pyrophosphate, 50 mM sodium fluoride, 40 mM b-gylcophosphate, 1 mM sodium vanadate, 100 mM phenylmethylsulfonylfluoride, 1 µg/mL leupeptin, and 5 µg/mL aprotinin) for 10 minutes on ice. Cells were then scraped and syringed with 22 gauge syringes. Cell lysates were boiled in 2x sample buffer (BioRad 1610737).

1205Lu cells were transduced with Halo-TRIM9ΔRING adenovirus and treated with 1 µg/ml Doxycycline (Sigma D9891) to induce expression. 24h post transduction, cells were transfected with VASP-GFP with Lipofectamine 2000 per manufacturer’s instructions. 16-18 h post-transduction, cells were washed with 5 mL of 1x PBS 3 times. Cells were crosslinked using 0.5 mM Pierce DSP (Thermo A35393) diluted in PBS for 25 min at RT on an orbital shaker. Cells were then washed with 5 ml 1x TBS. Cells were incubated in cytoplasmic extraction buffer (1 ml TBS, 1% NP-40, 0.1% SDS) for 25 min at RT on an orbital shaker. Lysates were collected and centrifuged at 4C for 5 min at 600 x g. Lysates were diluted to 1 mg/ml in IP buffer (containing 10% glycerol, 50 mM Tris pH 7.5, 200 mM NaCl, 2mM MgCl2, 1% NP-40, with 15 mM sodium pyrophosphate, 50 mM sodium fluoride, 40 mM b-gylcophosphate, 1 mM sodium vanadate, 100 mM phenylmethylsulfonylfluoride, 1 µg/mL leupeptin, and 5 µg/mL aprotinin). Protein concentrations were determined using a Bradford assay (BioRad 5000006) then normalized before IP. 1 mg of protein was rotated with 20 µL of GFP-TRAP beads (gtma, Proteintech) for 1 h at 4C. Beads were then isolated, washed with RIPA buffer 6 times for 10 min each, DSP crosslinking cleaved with 50mM DTT at 37 C for 30 min, and then boiled with 2x sample buffer.

Lysate samples and immunoprecipitants in sample buffer were loaded onto homemade 8 or 10% SDS-PAGE gels and run at 100V and subsequently transferred onto 45 µm nitrocellulose membrane (BIO-RAD 1620115) for 90 min (75 V). Membranes were blocked in 5% milk in 1x Tris-buffered saline (TBS) or 5% BSA (Roche 10735086001) in 1x TBS for 1 h at room temperature (RT). Blocked membranes were then incubated with primary antibody overnight in 2% milk or 2% BSA in TBS with 0.1% Tween-20 (TBST) on a rocker at 4C. Membranes were washed three times (5 min) in 1x RT TBST and subsequently incubated with secondary antibodies for 1 h on a rocker at RT, washed 3× 10min in 1x TBST, rinsed in PBS and imaged on an Odyssey (LI-COR Biosciences).

### Cell Spreading

Prior to plating, coverslips were cleaned using Harrick Plasma Cleaner (PDC-32G). 1205Lu were replated at 1.5×10^6^ on #1.5 coverslips in a 6 well plate. Two h after plating cells were fixed in 4% PFA PHEM fix for 10 min. Coverslips were then washed in 1x PBS three times prior to labeling with phalloidin (described in immunostaining section). Coverslips were imaged for phalloidin. Imaging and analysis were completed blinded. Phalloidin images a threshold was applied evenly across all images in ImageJ then quantified for cell area, perimeter, and intensity.

### Cell Blebbing

Gradient polyacrylamide gels with varying stiffness levels were generated (Hakeem et al., 2023). Briefly, 30 mm coverslips were washed with 100% ethanol, air-dried, and plasma cleaned (machine info) for 2–3 min. Coverslips were immersed in 0.5% 3-Aminopropyl triethoxysilane (APTES) for 20 min then washed with DI water 2 times for five min each. Coverslips were immersed in 0.5% Glutaraldehyde (Electron Microscopy Sciences, 16320) for 45 min then washed by immersing in water 3 times, 10 min each, and then airdried. Activated coverslips were stored in 4 C. A 12% acrylamide and 0.6% bis-acrylamide solution in Dulbecco’s PBS (DPBS) was prepared. A lithium phenyl-2,4,6-trimethylbenzoylphosphinate or LAP (TCI chemicals, L0290) solution (0.5% v/v) was made using the acrylamide mix in a glass vial (500 µL volume). The glass vial containing the mixture was covered in aluminum foil, protected from light exposure, and degassed in vacuum chamber for 15 min. The chemically activated 30 mm glass coverslip was placed in a 3D printed sample holding mold and a 15 µL drop of the LAP acrylamide mixture was placed in the center of the coverslip. A 12 mm glass coverslip was gently lowered over the drop to create a sandwich. The 3D-printed mold with a house-shaped void on the top surface was placed with the “house roof” pointing left and inserted into the path of light source. The bottom half of the house-shaped void was covered with an opaque paper mask. The sandwich was exposed for 3 min. At the 3 min mark, the paper slip was quickly and completely removed, and the sample was further exposed to light for another 45 s generating areas of the gel with different stiffness ~ 75 kPa and ~ 2 kPa. Light was quickly turned off, the top 12 mm coverslip was gently dislodged with fine forceps, and the gel was rinsed and stored in DPBS at 4 °C. Stored gels were gently washed thrice with DI water. The 1 mg/mL Sulfo-SANPAH (Thermo, 22589) solution was prepared in DI water from a concentrated 25 mg/mL stock in DMSO stored at 80 °C. The Sulfo-SANPAH solution was centrifuged at maximum speed (21,130 rcf) and passed through a 0.45 µm filter to remove particulate material. Filtered Sulfo-SANPAH solution was added to the gel surface and illuminated in a UV Stratalinker (Stratagene) oven for 30 min. Gel was then quickly transferred from the oven to tissue culture hood, rinsed 3 times with sterile water, and then incubated in 10 µg/mL laminin (Sigma #L2020) made in DPBS for 90 min at 37 °C. Excess laminin solution was aspirated, and gel was rinsed twice with DPBS and left immersed in DMEM prior to seeding and imaging of cells.

### Immunostaining

Prior to plating, coverslips were cleaned using Harrick Plasma Cleaner (PDC-32G). Cells plated on #1.5 coverslips were fixed in warm 4% PFA 1x PHEM (60mM PIPES 25mM HEPES, 10mM EGTA, 2mM MgCl2 pH 6.9) fix for 10 min at room temp. All the following steps are completed at RT. Coverslips were then washed in 1x PBS three times. Cells were permeabilized in 0.1% Triton X-100 in PBS (Fisher BP151) for 10 min, blocked in 10% normal donkey serum (Southern-Biotech 0030-01) in 1x PBS for 30 min, then washed with 1x PBS for 5 min. Coverslips were incubated in primary antibody in 1% normal donkey serum for 1 h at RT, washed with 1x PBS 3 times for 5 min each, then incubated with secondary antibodies and conjugated phalloidin-488 (Fisher A12379) and incubated at RT for 1 h then washed three times for 5 min each with 1x PBS prior to mounting in mounting media (a Tris, glycerol and n-propyl-gallate-based mounting medium made in house). All primary and secondary antibodies were diluted in 1% NDS-1x PBS. For all immunostainings, fluorescence intensities were normalized to the average of the mean intensity of conditions compared for each biological replicate.

### Actin Analysis

Lamellipodial width was measured manually using a 5 µm linescan drawn every 10 µm across the lamellipodia. The average length per cell was graphed. To analyze filopodial VASP localization and filopodial length and density, filopodia line scans were drawn manually from tip to base in the phalloidin channel in ImageJ. To attain VASP intensity at filopodia, the line scan drawn in the phalloidin channel was applied to the VASP channel. The perimeter of each cell via phalloidin was taken in ImageJ to calculate density. Filopodial width was quantified using line scans at the tip, middle, and base. The Full width half maxima (FWHM) was extracted from individual line scans per filopodia and averaged to have one width per filopodia. Filopodia width was quantified by taking the FWHM of the base, middle, and tip of a filopodia and taking the average. For VASP distribution analysis, to normalize filopodia length variations, each line intensity profile was binned into 20 bins, with bin 1 corresponding to the tip and bin 40 to the base. The average intensity value within each bin is assigned to that bin. The binned profiles were then pooled and averaged to create a heatmap for visualization (Ho et al., 2025). To quantify the thickness/bundling of stress fibers, maximum intensity projections were taken in ImageJ of phalloidin stained cells. The background was subtracted using rolling ball tool in ImageJ. Each ventral stress fiber had a central line scan drawn perpendicular to the fiber, the FWHM of each fiber was extracted.

### Focal Adhesion Analysis

1205Lu and WM266-4 cells were plated on 10 µg/ml FN (Corning 356008) for 3 h then subsequently fixed in 4% PFA 1x PHEM fix for 10 min at RT, washed, and immunolabeled for paxillin (1:200; anti-mouse BD Transduction Laboratories 610051), VASP (1:1000; anti rabbit VASP2010), and phalloidin-488 (1:400; Fisher A12379) as described in the immunostaining section. Coverslips were imaged using UP-lanApo 100×/1.5-NA DIC TIRF objective (Olympus). Prior to analysis, images were background subtracted using the rolling ball algorithm in ImageJ. For analysis of focal adhesion number, morphology and intensity, paxillin-positive adhesions were manually segmented using the freehand tool in ImageJ and saved as individual regions of interest (ROI). Area, mean gray value, integrated pixel intensity, and shape descriptors were extracted from segmentations.

### FRAP

VASP, VASP^KR^, paxillin and zyxin were imaged by TIRF illumination at an evanescent wave depth of 100 nm. Prebleach images were acquired at 10.000 s intervals for 5 frames with 50 ms exposure with 491 or 561 laser with 30% power. Using 405nm laser, a small ROI was placed on an individual focal adhesion, which was bleached for 10 ms with 100% power to 20-30% original fluorescence. Cells were then imaged for 400 frames at 0.300 ms intervals, then for 30 frames at 1s intervals with 50 ms exposures of 491 or 561 nm laser with 35% power.

To analyze FRAP experiments, all measurements were taken using a circle ROI within the bleach region with a diameter of 4.03 µm. The intensities of the FRAP ROI, three different focal adhesions were used to assess photofading (cell), and the background (bkgd) were measured. The following equation was used to correct for background fluorescence photofading, and loss of fluorescent material due to the bleach, and normalized to correct for differences in protein expression between cells (Day et al., 2012).

The normalized intensity of the FRAP ROI for each focal adhesion was fit in GraphPad Prism to a nonlinear regression (curve fit) exponential decay model (Y=(Y0 - Plateau)*exp(-K*X) + Plateau). The plateau value was extracted from the curve, and fluorescent recovery halftime was derived using: ln(2)/K.

### Collagen Gel Contraction

Collagen gels were generated by adding 500 µl of 10× MEM (Gibco 11430-030), 200 µl culture medium (DMEM and 10% FBS), and 270 µl of 7.5% sodium bicarbonate (Gibco 25080-094) to 3.75 ml of 3 mg/ml rat tail type-I collagen (Gibco A1048301). 765 µl of this mixture was added to a tube containing 1 million cells in 1.235 ml of DMEM + 10% FBS. Cells were quickly resuspended, and 400 µl of this mixture was added to wells in a 12 well plate and allowed to gel at 37°C with shaking every 15 min to prevent the sedimentation of cells. For controls, gels were created in different wells of the same 12 well plate without any cells. Technical replicates were plated. After 1 h, 1 ml of media was added to each well, and a P20 pipette tip was used to gently separate the gel from the well. Samples were placed in an incubator for 24 h before imaging plates using white light setting on ChemiDoc XRS system (Biorad). The area of each collagen gel was analyzed using ImageJ.

### Cell Migration

12-24 well glass bottom plates were coated with 10 µg/ml Laminin (Sigma #L2020), 10 µg/ml Fibronectin (Corning 356008), 1x PDL (Gibco A38904), or no coating overnight at 4C. Cells were allowed to fully spread after plating. Cells were imaged at 37 C, 5% CO_2_, with a Keyence PlanFluor 10x/0.45-NA Ph1 objective for 16 h at 3-8 min intervals. The manual tracking plugin on ImageJ was used to track single cells based on nuclear location. Cell tracks were terminated when cells collided, divided, died, or migrated outside of the field of view. Cells had to be tracked for a minimum of 30 minutes to be included in analysis. The ImageJ Chemotaxis Tool plugin was used to extract velocity and directional persistence from raw data.

### Transwell Invasion

In a 24 well plate, 0.8 µm pore transwells (Greiner bioone #662638) were coated with 30 µg of Geltrex (Gibco A40000469) in 100 µl of cold 1x PBS and placed at 37 °C to solidify for 1 h. After 1 h, 650 µl of DMEM supplemented with 5% FBS, to serve as a chemoattractant, was added to the bottom of the well. Cells were seeded at 25,000 cells/200 µl in DMEM with 0% FBS and returned to 37 °C for 20 h. After 20 h, media was removed from the top of the well, cells on top of transwell were removed using QTips, and subsequently fixed in in 4% PFA PHEM fix for 10 min. Transwells were then washed in 1x PBS three times prior, permeabilized with 0.3% Triton x-100 in PBS (Fisher BP151) for 10 min, stained with 300nM DAPI for 1 min, and washed with 1x PBS 3 times. Transwell membranes were removed from the transwell insert with a scalpel and mounted in mounting media. Transwell membranes and the corresponding well were imaged on Keyence PlanFluor 10x/0.45-NA Ph1 objective to count cells that had migrated through the transwell. The fraction of cells invaded is reported.

### Zymography

80% confluent 6 cm dishes were rinsed with PBS and covered with 4 mL of serum-free DMEM. After 24 h, 3.5 mL of this conditioned media was removed and concentrated to 250 µL using Amicon Ultra-15 Centrifugal Filters. Conditioned media volumes were normalized according to cell count from the dishes of origin and incubated with 2 × SDS loading buffer without 2-mercaptoethanol for 20 min at 37 °C before being loaded into a 10% SDS gel containing 0.1% porcine gelatin. Gels were run at 200 V packed on ice until the 37kd marker ran off of the gel, then washed four times for 30 min each at room temperature with wash buffer (25 mM Tris pH 7.6, 5 mM CaCl_2_, 3 mM NaN_3_, 0.25 nM ZnCl_2_, and 1.25% Triton-X100 in water). The gel was then washed with ddH2O for 30 min before incubation for 36 h at 37°C in incubation buffer composed of 25 mM Tris pH 7.6, 5 mM CaCl_2_, 3 mM NaN_3_ and 0.375 M NaCl in water. The gel was then quickly rinsed in distilled water before staining with 0.1% Coomassie Blue solution for 1 h. After destaining for about 30 min, the gel was imaged using an Odyssey (LI-COR) Biosciences) Imaging System. Band intensity was quantified using Licor Studio Lite and graphed in GraphPad Prism.

### Cell proliferation assay

Cells were trypsinized, counted using the TC20 Automated Cell Counter (Biorad), and 1 ml of culture media containing .05 × 10^6^ cells were plated into each well of a 12-well plate (day 0). After 24 h (day 1), cells were trypsinized and counted. This was repeated for the following 5 d. Three independent biological replicates were performed with technical duplicates.

### MTT cell proliferation assay

Cell proliferation was evaluated using an MTT (3-(4,5-Dimethylthiazol-2-yl)-2,5-Diphenyltetrazolium Bromide, Sigma-Aldrich, Cat# 475989) assay per manufacturer protocol. Cells were seeded in 96-well plates at a density of 3,000 cells per well in 100 µL of culture medium. Cell viability was measured at 1, 3, and 6 days post-plating in triplicate. At each time point, culture medium was removed and replaced with 100 µL of MTT working solution prepared by diluting MTT 1:10 in serum-free DMEM to a final concentration of 500 µg/mL. Cells were incubated for 1 hour at 37 °C to allow formation of formazan crystals. The MTT solution was then removed and the crystals were solubilized in 200 µL dimethyl sulfoxide (DMSO) by pipetting until fully dissolved. Absorbance was measured at 570 nm using a microplate reader.

### Tumor cell proliferation in vivo

To evaluate differences in tumor cell proliferation dynamics and tumor depth growth dynamics between *PBT:Trim9*^*+/+*^ and *PBT:Trim9*^*f/f*^ groups, a multi-level analytical approach was employed. Cell density measurements were collected using two photon imaging at days 1, 5, 8, 13, and 15 post-induction (n = 4 replicates per group). Tumor depth measurements were collected at weeks 4 through 13 post-induction. Not all replicates were measured at every timepoint; missing values were handled by smoothing each replicate over its available observations only.

Growth trajectories were modeled using Functional Data Analysis (FDA), implemented via the fda package in R (v4.5.2). Individual replicate measurements were smoothed using cubic B-spline basis functions (order = 4, nbasis = 5) with a roughness penalty parameter (lambda = 0.1) to generate continuous functional representations of each growth curve. Each replicate was smoothed independently over its available timepoints to accommodate missing data. Mean curves and first derivatives (instantaneous growth rates) were estimated for each group across the full observation period (days 1–15).

To formally test for differences in overall curve shape between groups, a permutation test (n = 999 permutations) was performed using the mean squared difference between group mean curves as the test statistic. Group labels were permuted across all replicates to generate a null distribution, and a p-value was calculated as the proportion of permuted statistics equal to or exceeding the observed statistic. Additionally, the slope of cell density change between days 5 and 8 was calculated for each replicate as density / time, and group differences in slope were compared using a two-sample Welch’s t-test. All statistical analyses were performed in R (v4.5.2). A significance threshold of α = 0.05 was applied; results with p < 0.10 were considered to represent statistical trends.

To assess total proliferative output, area under the curve (AUC) was calculated for each replicate using the trapezoidal method (DescTools package), and group differences in AUC were tested using a two-sample Welch’s t-test.

Given the visual divergence in growth trajectories observed after day 5, a focused analysis of the late-phase growth rate was performed. The slope of cell density change between days 5 and 8 was calculated for each replicate as density / time, and group differences in slope were compared using a two-sample Welch’s t-test. All statistical analyses were performed in R (v4.5.2). A significance threshold of α = 0.05 was applied; results with p < 0.10 were considered to represent statistical trends.

### Cell Shape Analysis in vivo

A z-stack of 20x multiphoton image of tdTomato 5 days post tumor induction were acquired, and a max projection was made. Image analysis was conducted blinded. Images were first smoothened to reduce noise and lessen pixelation. Using the IsoData threshold in FIJI, individual cells were selected with the magic wand tool to measure parameters of cell shape.

### Exponential Growth Curve Analysis of Tumor Depth

Tumor depth over time was analyzed by fitting an exponential growth model in GraphPad Prism. For each genotype, tumor growth measurements were plotted as a function of time and fit using Prism’s nonlinear regression exponential growth equation: Y=Y_0_ x e^kt^ where *Y* represents the measured signal, Y_0_ is the starting value, *k* is the exponential growth rate constant, and *t* represents time.

To determine whether the growth dynamics differed between genotypes, curves were compared using Prism’s extra sum-of-squares F test within the nonlinear regression framework. Two models were evaluated: (1) a global model in which both genotypes were constrained to share the same growth parameters and (2) a separate-curves model in which parameters were allowed to vary independently for each genotype. The models were compared using an F test (α = 0.05) to determine whether allowing separate curves significantly improved the fit.

### Macro-metastasis identification

Mice were sacrificed between 13-16 weeks via CO_2_ euthanasia and cervical dislocation according to UNC’s Institutional Animal Care and Use Committee guidelines. Organs were harvested, placed in sterile PBS and imaged with an Olympus MVX10 macroscope system. TdTomato + macro-metastases were identified using fluorescence and appropriate filter sets in conjunction with Metamorph software.

### Microscope Descriptions

FRAP and TRIM9 localization experiments were performed on an Olympus IX83-ZDC2 equipped with a cMOS camera (Orca-Fusion, Hamamatsu), Xenon light source (Sutter), and the following objective lenses: a UPlanApo 100×/1.51 NA DIC TIRF objective (Olympus), a UAPON 100×/1.49-NA DIC TIRF objective (Olympus using Cellsens software (Evident) acquisition software.

Cells for live imaging were maintained in culture medium and at 37°C and 5% CO_2_ with a Tokai-Hit on-stage incubator. Random cell migration assays, WM266-4 blebbing assays, spreading assays were conducted on Keyence BZ-X800 All-in-one Fluorescence Microscope using a Keyence PlanFluor 10x/0.45-NA Ph1 objective, a Keyence PlanFluor 10x/0.45-NA Ph1 objective, Keyence PlanFluor 40x/0.60-NA Ph2 fluor objective, respectively. Cell blebbing assays for 1205Lu cells were imaged using a Olympus VivaView FL microscope at 37 °C and 5% CO_2_ with a 20× objective at 10 min intervals for 30-36 h.

Multiphoton imaging experiments were conducted on a Olympus FV1000 MPE confocal and Multiphoton system on a Olympus BX61WI stand with remote focus module, equipped with a 20x/1.0 water XLUMPlanFL objective, a Prior Z Deck motorized stage with ProScan III controller, Spectra-Physics Insight DeepSee laser tuned to 910nm and controlled by Fluoview software FV10-ASW 4.1. Gross mouse ear images were acquired on a Leica MZ16FA motorized fluorescence stereo microscope with 0.63X Planapochromatic lens with a DsRed filter set and Micropublisher 5.0 (Q-Imaging) camera, controlled by MetaMorph software (version 7.10.3.279, Molecular Devices).

### Statistics

Statistical analyses were completed in Graphpad Prism10 (RRID:SCR_002798). Plots were made in Superplots (Goedhart, 2021). A minimum of three biological experiments were performed for data performed with statistics. The number of replicates and statistical tests are indicated in figure legends. Unless indicated otherwise, data distributions were normal. For normally distributed data, two-tailed *t* tests were performed when two groups were compared, or one-way ANOVA or two-way (Bonferroni’s or Dunnet’s post hoc multiple comparisons test). For data that were not distributed normally, Mann-Whitney or Kolmogorov-Smirnov tests were used. Statistical differences were determined by an α= 0.05.

## Data Availability Statement

The data generated in this study are available from the corresponding author upon request.

## Acknowledgements

We thank Janee Cadlett-Jette and Natallia Riddick for assistance with mouse husbandry. We thank Dr. Nancy Thomas for thoughtful conversations and generous sharing of data. We thank Wendy Salmon in the Hooker Imaging Core for her expertise and training. We thank Jadiel A. Aviles-Lukasik for generating a python script to extract FWHM based analysis. We thank Elliot Evans for his assistance in genotyping. We thank Max Hockenberry for guidance on actin bundle analysis, Bhagawat C. Subramanian for assistance with flow cytometry and tumor induction guidance, Mark Hazelbaker for helping find antibodies, and Ian C. McCabe for training in the transwell invasion assay.

This work was supported by National Institutes of Health grants R35GM135160 (S.L.G.), R35GM130312 (J.E.B.), a Developmental Pilot Project award from the UNC Lineberger Comprehensive Cancer Center (S.L.G.). K.L. was supported by a National Cancer Institute F99/K00 fellowship F99CA294269, Howard Hughes Medical Institute Gilliam Fellowship GT15781, National Institutes of Health T32GM119999-05, National Institutes of Health R25GM05536. Microscopy performed at the UNC Hooker Imaging Core Facility was supported in part by P30 CA016086 Cancer Center Core Support Grant to the UNC Lineberger Comprehensive Cancer Center.

## Author contributions

Conceptualization: K.L., J.E.B., S.L.G.; Methodology: K.L., C.T.H, S.R.; Formal analysis: K.L., C.T.H., J.B., A.B.S.; Investigation: K.L., A.B.S., G.B.P., C.T.H, M.L.; Resources: S.L.G.; Writing - original draft: K.L, S.L.G.; Writing - review editing: K.L., C.T.H., A.B.S, M.L, G.B.P., J.B., S.R., J.E.B., S.L.G.; Visualization: K.L., S.L.G.; Supervision: S.L.G.; Project administration: S.L.G.; Funding acquisition: S.L.G., K.L.

## Supplementary Information

**Supplemental Video 1. TRIM9 promotes bleb-like states on soft substrates**. DIC timelapse imaging of 1205Lu *TRIM9*^*+/+*^ and *TRIM9*^−/−^ cells on 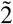 kPa polyacrylamide hydrogels.

**Supplemental Video 2. TRIM9 localizes to VASP+ structures**. TIRF timelapse imaging of 1205Lu cells expressing mCherry-TRIM9DRING and GFP-VASP.

**Supplemental Video 3. FRAP of VASP at focal adhesions**. TIRF-FRAP timelapse imaging of 1205Lu *TRIM9*^*+/+*^ and *TRIM9*^−/−^ cells expressing GFP-VASP.

**Supplemental Video 4. Random migration**. Phase contrast timelapse imaging of 1205Lu *TRIM9*^*+/+*^ and *TRIM9*^−/−^ cells migration on laminin coated coverglass.

**Supplemental Video 5. Images of *PBT-Trim9+/+* primary tumors**. Transmitted light images of tumors at 16 weeks post tumor induction.

**Supplemental Video 6. Images of PBT*-Trim9f/f* primary tumors**. Transmitted light images of tumors at 16 weeks post tumor induction.

**Supplementary Fig. 1.**
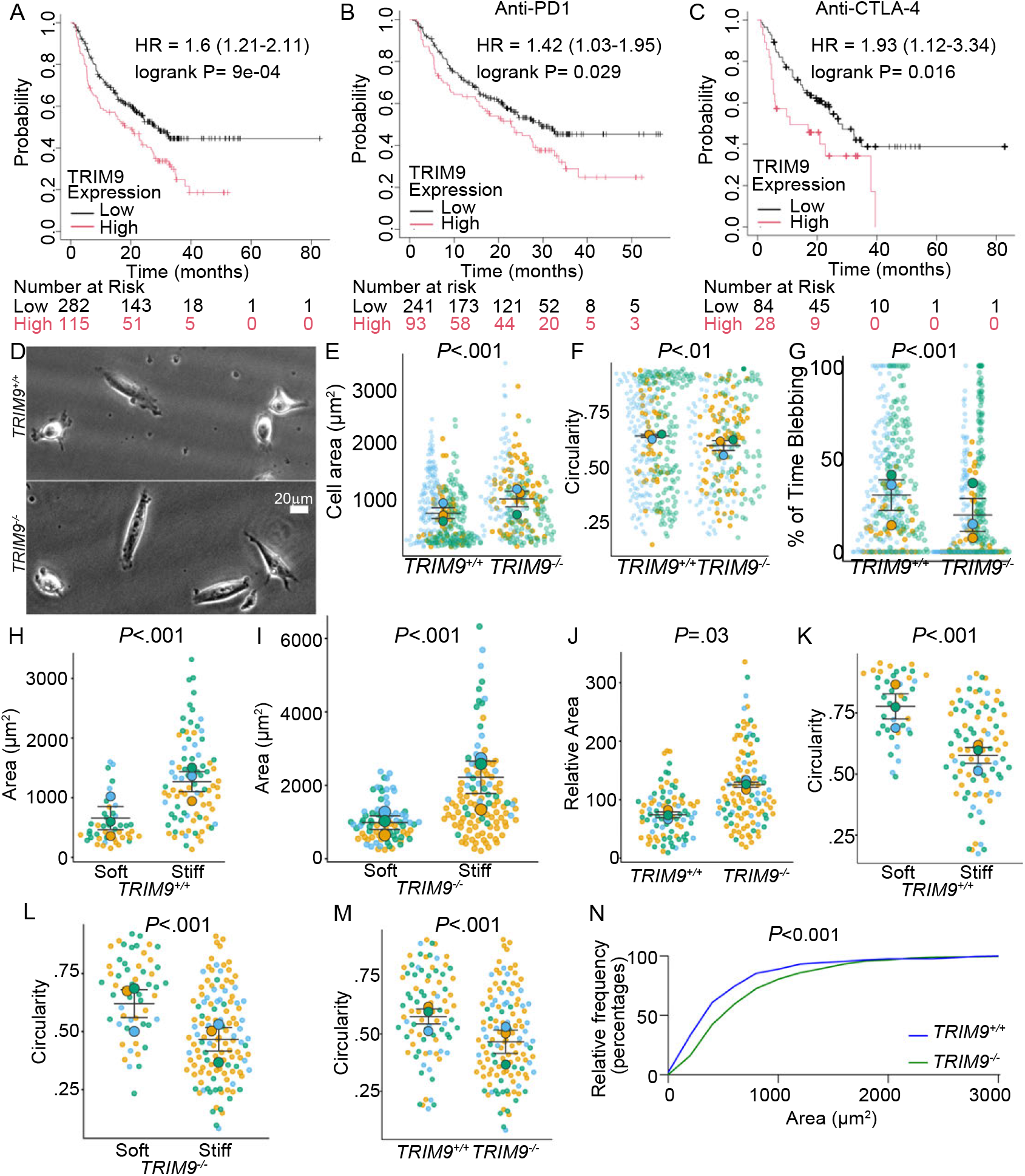
TRIM9 is expressed in human melanoma and alters cell morphology. (**A-C**) Kaplan-Meier analysis of overall survival stratified by TRIM9 mRNA expression in patients with melanoma either untreated (A), anti-PD1-treated (B) or anti-CTLA-4 treated (C). Patients were separated into high and low expression groups based on median TRIM9 expression. Survival differences were evaluated using the log-rank test. Hazard ratio (HR) and 95% confidence interval (CI) were calculated using Cox proportional hazards regression. (**D**) WM266-4 cells on laminin coated coverglass. (**E-F**) Loss of TRIM9 in WM266-4 increased cell area and circularity, (*P* <.001, *P* <.01, Mann Whitney test), n = 3 independent biological replicates; n 50 cells per replicate. (**G**) Loss of TRIM9 in WM266-4 cells reduced % of time in bleb-like morphology (*P* <.001, Mann-Whitney test), n = 3 independent biological replicates, each color is a replicate; n ≥ 37 cells per replicate, each dot is a cell. (**H-I**) Cell area of 1205Lu cells area reduced on soft (2kPa) relative to stiff hydrogel substrates (75kPa) (*P*<.001, two-tailed *t*-test), n = 3 independent biological replicates, each color is a replicate; n ≥ 7 cells per genotype per replicate, each dot is a cell. (**J**) Relative cell area of 1205Lu cells on stiff gels indicated increased cell spreading due to loss of TRIM9 (*P* =0.03, Two-tailed paired *t*-test), n = 3 independent biological replicates, each color is a replicate; n 12 cells per genotype per replicate. (**K-L**) Circularity of 1205Lu cells on soft (2kPa) vs stiff hydrogel substrates (75kPa); both genotypes were more circular on soft substrates (*P*<.001, two-tailed *t*-test), n = 3 independent biological replicates; n 7 cells per genotype per replicate. (**M**) Cell circularity of 1205Lu cells; loss of TRIM9 reduced cell circularity on stiff substrates (*P*<.001, two-tailed *t-*test), n = 3 independent biological replicates, each color is a replicate; n ≥ 12 cells per genotype per replicate, each dot is a cell. (**N**) Frequency distribution of 1205Lu cell area after two hours of spreading on coverglass, (*P*<0.001, Kolmogorov-Smirnov test).

**Supplementary Fig. 2.**
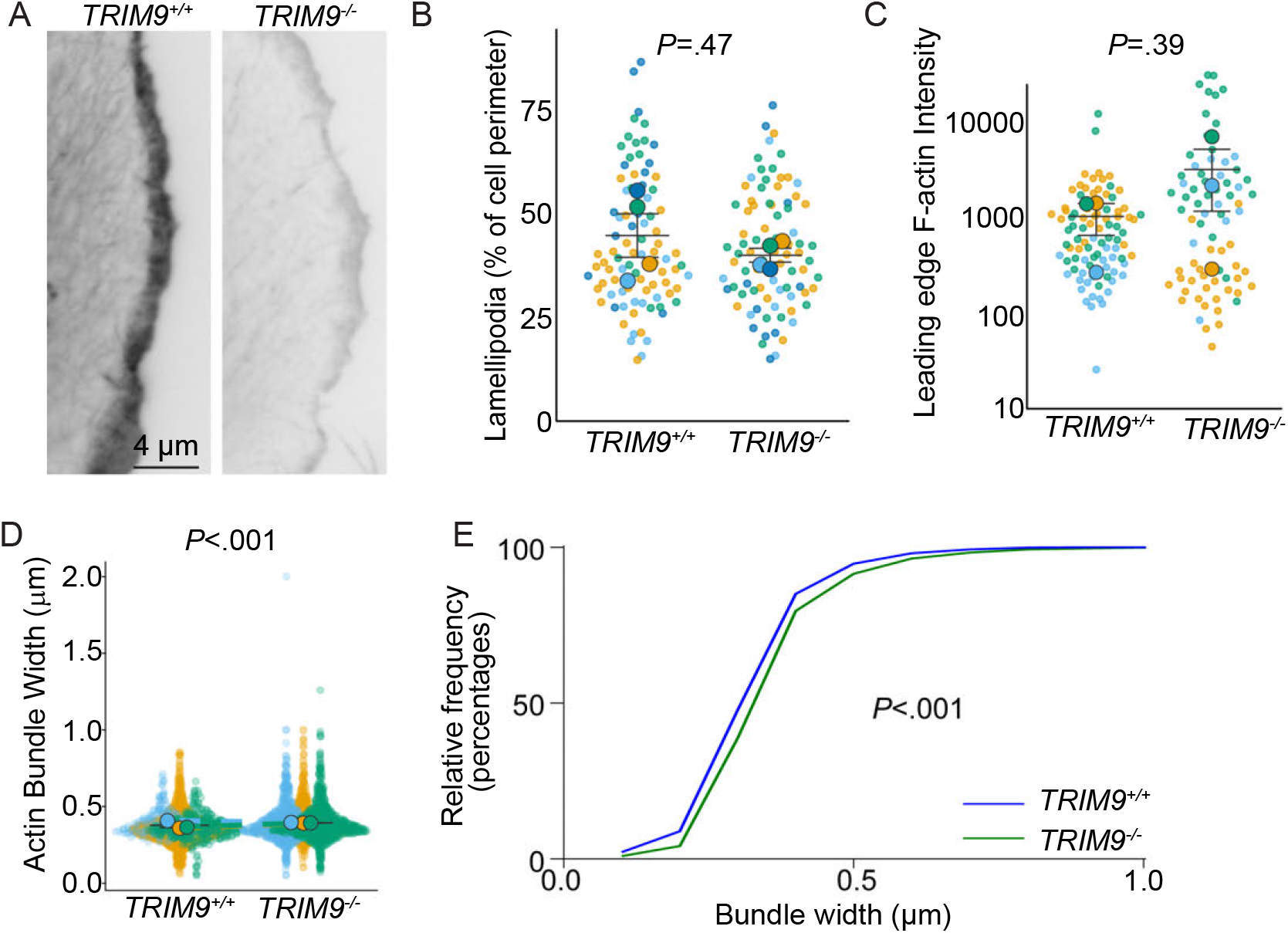
Loss of TRIM9 alters F-actin organization. (**A**) F-actin labeled 1205Lu cells with lamellipodial protrusions, but do not have filopodia. (**B**) Lamellipodial length respective to cell perimeter is not altered due to the loss of TRIM9 (*P* =0.47, Two-tailed *t*-test), n = 4 independent replicates with n ≥ 11 cells per genotype per replicate. (**C**) F-actin intensity at the leading edge of 1205Lu cells; loss of TRIM9 did not alter actin at edge of cell (*P* =.387, Two-tailed *t*-test), n = 3 independent replicates; n ≥ 13 cells per genotype per replicate. (**D-E**) F-actin bundle width in 1205Lu cells; loss of TRIM9 increased bundle width, (*P* < .001 with both Mann-Whitney and Kolmogorov-Smirnov test), n = 3 independent replicates; n ≥ 152 bundles per genotype per replicate.

**Supplementary Fig. 4.**
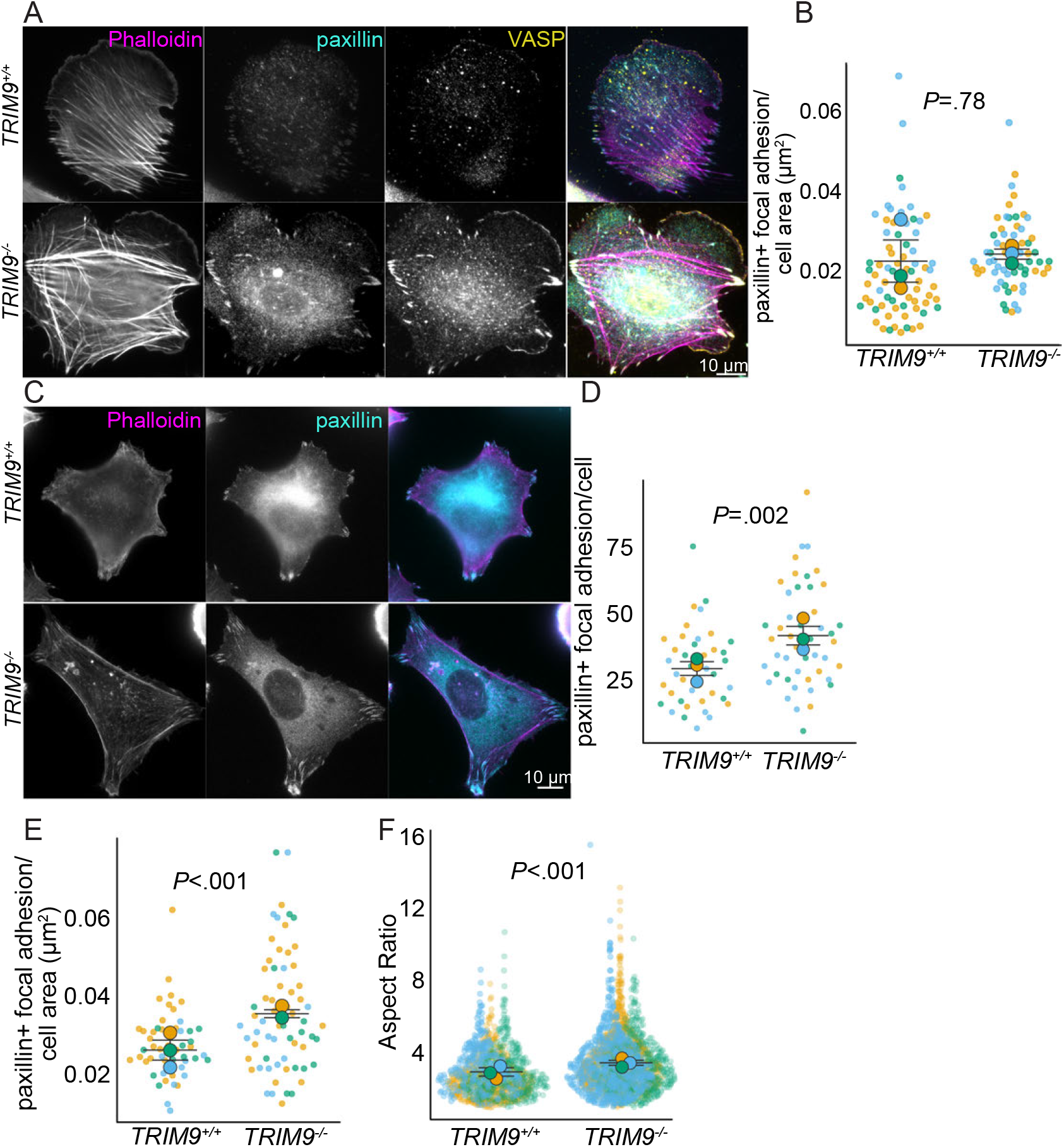
Loss of TRIM9 alters VASP localization to focal adhesions, focal adhesion number, and morphology. (**A**) Same representative images from Figure 4A, of 1205Lu cells, brightness and contrast adjusted differently to visualize focal adhesions in the *TRIM9*^*+/+*.^ cells. (**B**) Focal adhesion density in 1205Lu cells; loss of TRIM9 did not alter focal adhesion when normalized to area (*P* =.78, two tailed *t*-test) n = 3 independent biological replicates, each color is a replicate; n ≥ 7 cells per genotype per replicate, each dot is a cell. **(C)** Phalloidin and paxillin stained WM266-4 cells. stained WM266-4 cells. (**D)** Focal adhesion number in WM266-4 cells; loss of TRIM9 increased focal adhesions per cell (*P* =.002, two-tailed *t-*test), n = 3 independent biological replicates; n ≥ 11 cells per genotype per replicate. (**E**) Focal adhesion density in WM266-4 cells; loss of TRIM9 increased the density of focal adhesions (*P* <.001, two-tailed *t*-test), n = 3 independent biological replicates, each color is a replicate; n 11 cells per genotype per replicate, each dot is a cell. (**F**) Loss of TRIM9 in WM266-4 cells increased focal adhesion aspect ratio (width: height ratio) (*P* <.001, Kolmogorov–Smirnov test), n = 3 independent biological replicates, each color is a replicate; n ≥ 275 focal adhesions per genotype per replicate, each dot represents focal adhesion.

**Supplementary Fig. 5.**
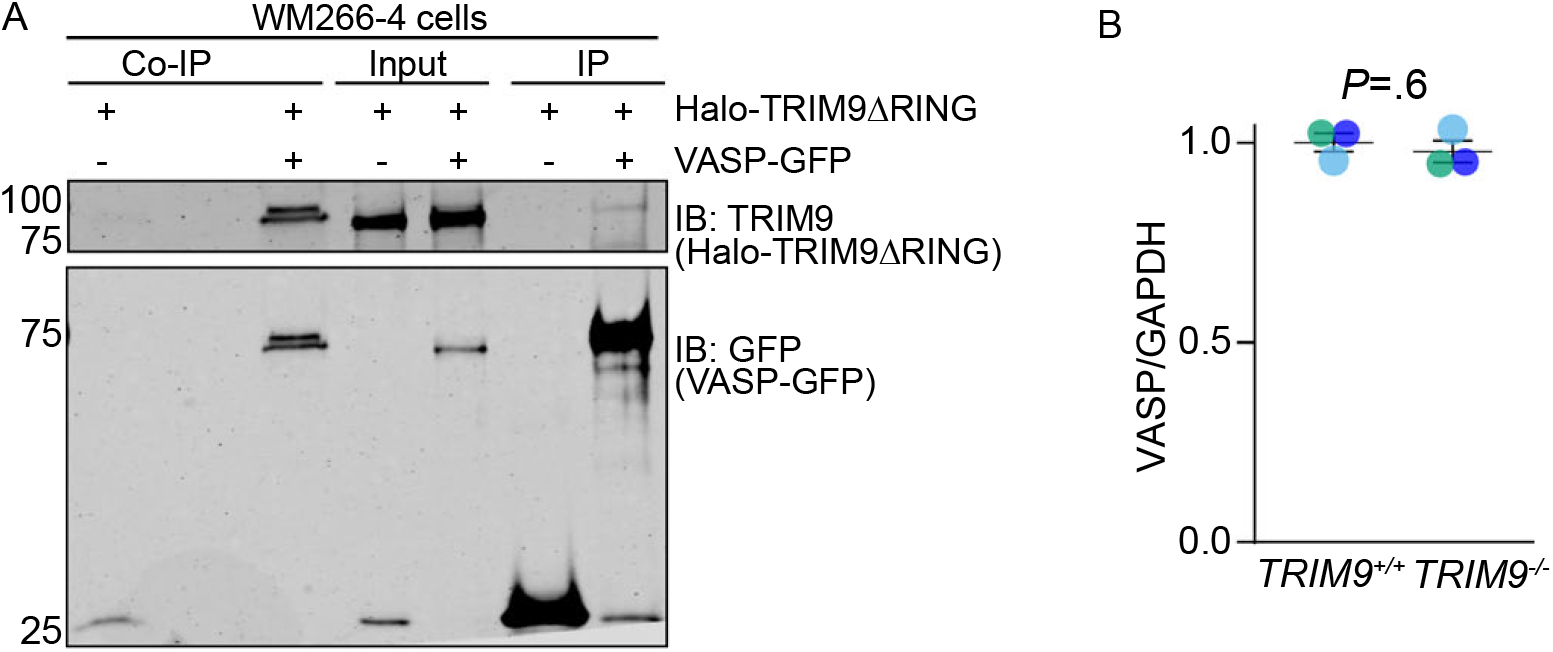
TRIM9 interacts with VASP in human melanoma and increases VASP phosphorylation. (**A**) Representative co-immunoprecipitation assays (Co-IP) from Dithiobis (succinimidyl propionate) (DSP)-crosslinked lysates from WM266-4 cells transfected with VASP-GFP or GFP and transduced with an adenoviral HALO-TRIM9ΔRING demonstrate an interaction between VASP and TRIM9 (n = 3 independent replicates). (**B**) VASP levels in 1205Lu cells normalized to GAPDH levels indicate loss of TRIM9 did not affect SDS-soluble VASP level (*P*=.6, paired two-tailed *t*-test), n = 3 independent biological replicates, each color is a replicate. This quantification corresponds to representative western blot in figure 5B.

**Supplementary Fig. 6.**
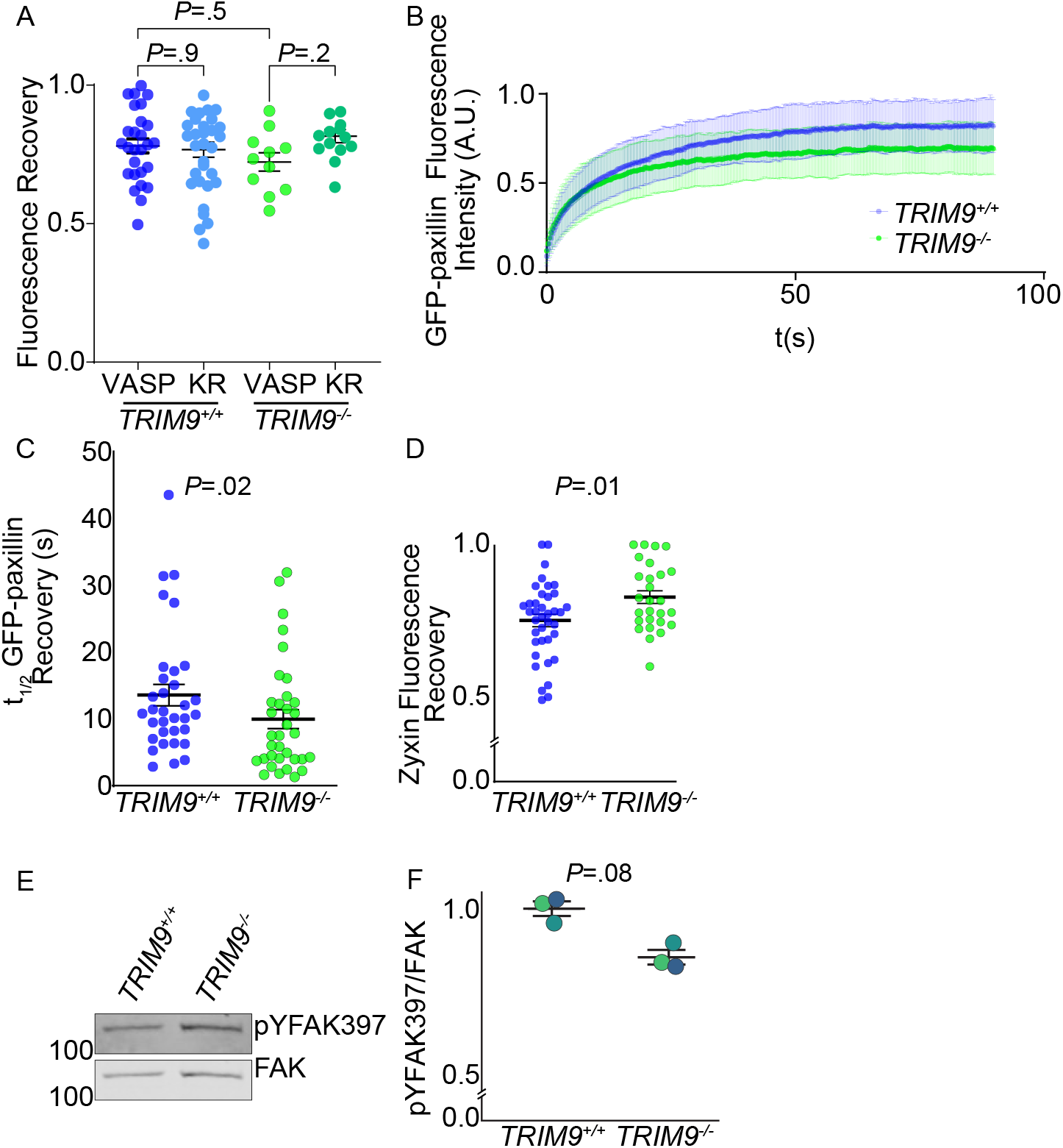
TRIM9 regulates VASP, zyxin, and paxillin dynamics at focal adhesions. (A) Fluorescence Recovery of VASP-GFP and VASPKR-GFP after photobleaching, P values from One-way ANOVA with Bonferroni’s multiple comparisons test), n = 3 biological replicates; n ≥ 25 cells. (B) A normalized fluorescence intensity of GFP paxillin after photobleaching, n = 3 biological replicates; n ≥ 33 cells. (C) t1/2 kinetics of GFP-paxillin in 1205Lu cells; loss of TRIM9 increases paxillin focal adhesion recovery rates (P =0.02, two-tailed t-test on log transformed data), n = 3 biological replicates; n ≥ 33 cells. (D) Fluorescence recovery of zyxin-mCherry in 1205Lu cells; faster zyxin recovery occurred in TRIM9-−/− cells (P=0.01, two-tailed t-test), n = 3 biological replicates; n ≥ 25 cells. (E-F) Immunoblot of pYFAK at Tyrosine 397 normalized to total FAK; loss of TRIM9 did not perturb pYFAK397 levels, (P =0.08, two-tailed paired t-test), n =3 independent biological replicates.

**Supplementary Fig. 7.**
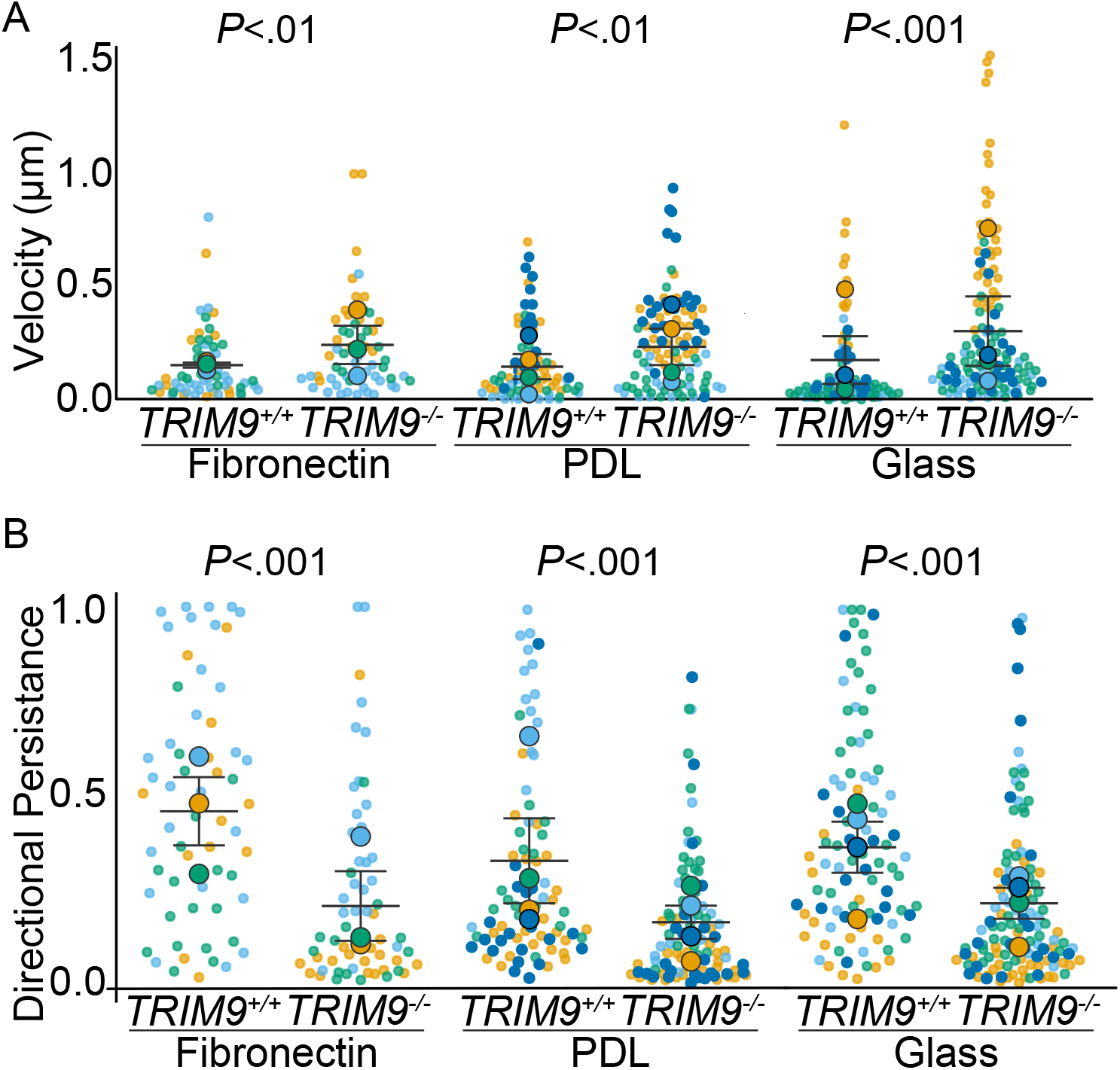
TRIM9 negatively regulates cell contractility, migration, and invasion in vitro. (**A-B**) 1205Lu cells plated on coverslips coated with 10 µg/ml fibronectin, PDL, or plasma-treated coverslips for 6 hours and imaged for 18 hours using phase contrast time lapse imaging. (**A**) Instantaneous velocity of 1205Lu cells *TRIM9*^−/−^ cells was increased compared to *TRIM9*^*+/+*^ cells on fibronectin (*P*<.01, two-tailed *t*-test on log transformed data), 1xPDL, (*P*<.01, sqrt transformed data), and plasma-treated coverslips (*P*<.001, two-tailed *t*-test on log transformed data).(**B**) The directional persistence of 1205Lu *TRIM9*^−/−^ cells was increased compared to *TRIM9*^*+/+*^ cells on fibronectin (*P*<.001, two-tailed *t*-test on log transformed data), 1xPDL, (*P*<.001, two-tailed *t*-test on log transformed data), and plasma-treated coverslips (*P* <.001 KS test).

**Supplementary Fig. 8.**
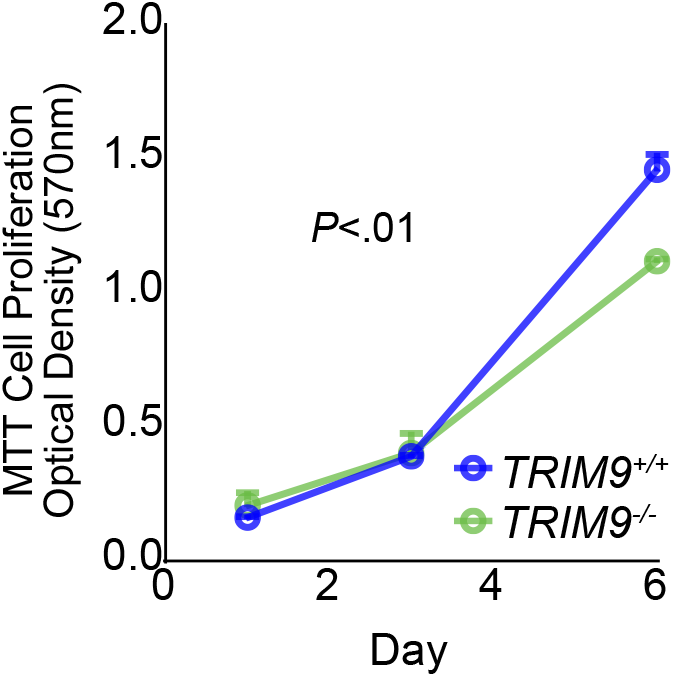
Loss of TRIM9 reduces proliferation. MTT assay evaluating cell proliferation; loss of TRIM9 reduced cell proliferation (*P*<.01, two-way ANOVA), n=3 independent biological replicates with 3 technical replicates.

**Supplementary Fig. 9.**
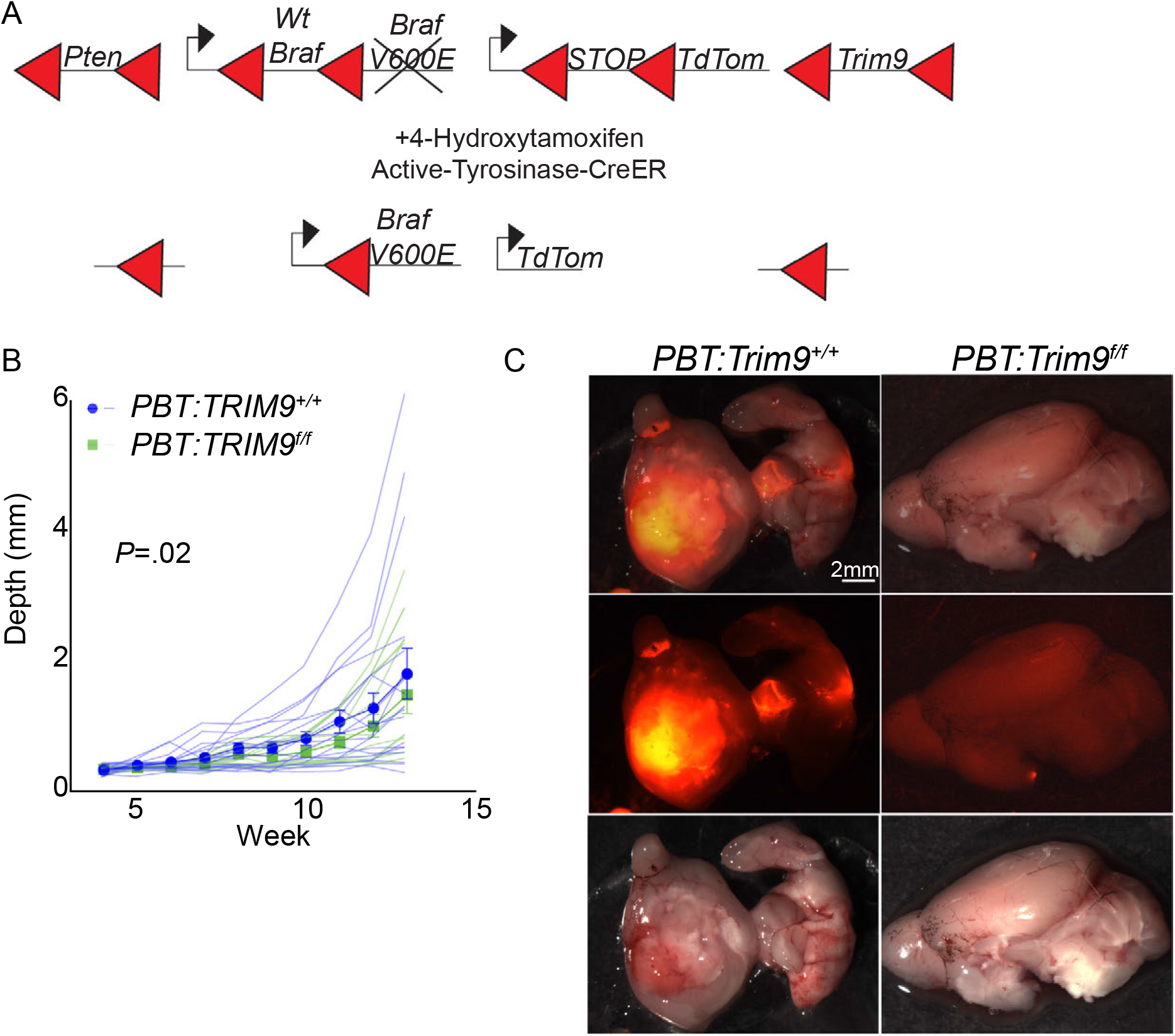
TRIM9 promotes tumor growth and alters metastasis in vivo. (**A**) Schematic of *PBT-Trim9*^*f/f*^ GEM recombination. Upon 4-HT application, CreER is expressed under the melanocyte-specific Tyrosinase promoter, excising the DNA between LoxP sites (red triangles), resulting in deletion of *Pten, wildtype Braf*, and *Trim9*, and expression of a constitutively active *BrafV600E* mutant and *TdTomato* in melanocytes. (**B**) Quantification of tumor depth over a 13-week period, as seen in Fig9G, with individual tumor depths measurements represented in light blue or light green. n= 18 *PBT-Trim9*^*+/+*^, 13 *PBT-Trim9*^*f/f*^ mice. (**C**) Macro-metastases in the brain of *PBT-Trim9*^*+/+*^ and *PBT-Trim9*^*f/f*^ mice.

